# Physics and physiology determine strategies of bacterial investment in flagellar motility

**DOI:** 10.1101/2024.03.30.587422

**Authors:** Irina Lisevich, Remy Colin, Hao Yuan Yang, Bin Ni, Victor Sourjik

## Abstract

Regulatory strategies that allow microorganisms to balance their investment of limited resources in different physiological functions remain poorly understood, particularly for numerous cellular functions that are not directly required for growth. Here, we investigate the allocation of resources to flagellar swimming, the most prominent and costly behavior in bacteria that is not directly required for growth. We show that the dependence of motile behavior on gene expression in *Escherichia coli* is determined by the hydrodynamics of propulsion, which limits the ability of bacteria to increase their swimming by synthesizing more than a critical number of flagellar filaments. Together with the fitness cost of flagellar biosynthesis, this defines the physiologically relevant range of investment in motility. Gene expression in all *E. coli* isolates tested falls within this range, with many strains maximizing motility under nutrient-rich conditions, particularly when grown on a porous medium. The hydrodynamics of swimming may further explain the bet-hedging behavior observed at low levels of motility gene expression.

## Introduction

Microorganisms, like all living systems, must achieve multiple physiological objectives that may change when encountering new environments. To perform successfully, microorganisms have therefore evolved numerous regulatory mechanisms responsible for allocating limited resources to specific physiological functions^1,2^. Bacteria, including *Escherichia coli* have become convenient models to address this fundamental resource allocation problem^3^, with a primary focus on proteome partitioning^4–7^. To allocate their proteomic resources into protein biosynthesis as a function of growth rate, bacteria appear to obey linear rules known as growth laws^4,5^: the fraction of the proteome responsible for biomass production expands with growth rate, whereas the fraction responsible for nutrient uptake and catabolism decreases with growth rate. This leads to the negative linear relation between the expression of carbon catabolic genes and growth rate, known as the C-line^5^, which has been proposed to maximize growth. However, although growth maximization is an important research allocation strategy^4–6,8^, it is not always the case^9,10^ and cells may instead prioritize other targets such as energy yield or stress response^11,12^. Furthermore, while previous studies have mostly focused on the optimized expression of catabolic^4,5,8,10^, anabolic^5,6^ or ribosomal^2,5,6^ genes, microbial strategies for resource allocation to multiple functions not directly required for growth remain unclear^13^.

The most prominent example of such a costly physiological function is swimming motility. Motile bacteria are propelled by the rotation of long helical flagellar filaments powered by a motor that is typically proton-driven^14^. Motility enables bacteria to follow spatial gradients of nutrients or harmful chemicals sensed by the chemotaxis signaling pathway^15,16^. Motility consumes several percent of total cellular resources in *E. coli* and other bacteria^17–19^, primarily due to the protein budget required for the biosynthesis of flagella^20,21^. Consistent with this high cost, several studies have observed a trade-off between growth and motility in *E. coli*^21–24^. However, the exact dependence of this trade-off on the absolute level of resource allocation to swimming motility remains uninvestigated.

Interestingly, the flagellar regulon in *E. coli* is controlled by catabolite repression^25^, such that flagellar gene expression increases in minimal medium during growth on poor carbon sources in accordance with the C-line^7,21^. The physiological relevance of such an investment strategy remains debated. One proposed explanation is that it ensures an anticipatory allocation of resources towards motility, in proportion to the potential benefit of finding additional nutrient sources via chemotaxis, which is higher in nutrient-poor environments^21^. Alternatively, it has been suggested that the number of flagella is tuned to match growth rate-dependent changes in cell size^23^.

In this study, we quantified the relation between the expression of motility genes and motile behavior, as well as the impact of motility on the growth fitness of *E. coli*. We demonstrate that major limitations on resource investment in motility, at both high and low levels of gene expression, arise from hydrodynamic constraints on bacterial swimming. Together with the fitness cost of flagellar synthesis and operation, this creates the physiologically relevant range within which the expression level of motility genes can vary depending on the conditions. We observe that within this range, *E. coli* follows different strategies of resource allocation towards motility depending on the medium, growth rate and isolate.

## Results

### Native regulation of motility genes in nutrient-rich medium maximizes swimming while limiting the cost of expression

To investigate how motility and growth depend on the expression of flagellar genes, we engineered a derivative of *E. coli* K-12 strain MG1655 with titratable expression of the *flhDC* operon that encodes the master activator of the entire flagellar regulon (Fig. 1a, Supplementary Table 1 and Methods). Expression of the flagellar regulon at different levels of *Ptac*-*flhDC* induction was quantified using a fluorescent reporter for flagellin (*fliC* gene) promoter activity (P*_fliC_*), which was previously shown to efficiently report the production of flagella in *E. coli*^20,21,26^. Reporter activity was measured using either a plate reader to follow changes in in the mean expression over time (Extended Data Fig. 1a, b), or flow cytometry to determine the distribution of single-cell expression levels within the cell population at a defined time point in mid-exponential phase (Fig. 1b). We confirmed that both readouts yielded similar results for *E. coli* cultures grown in nutrient-rich tryptone broth (TB) medium, with native MG1655 (wild-type; MG1655 *WT*) expression falling at an intermediate level within the range covered by the inducible *Ptac* strain (Fig. 1c and Extended Data Fig. 1b).

**Fig. 1.**
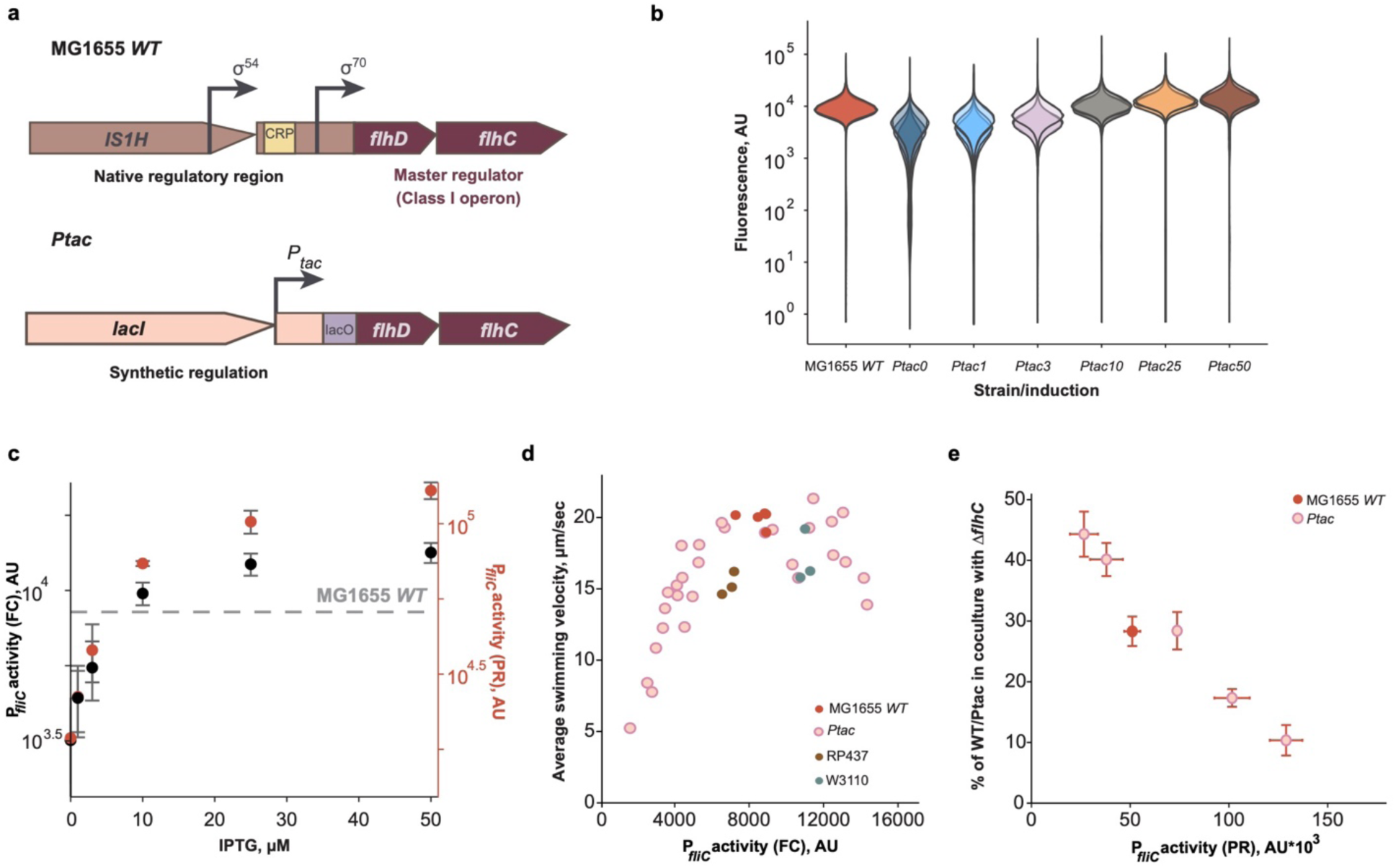
Dependence of motility and its cost on the expression of *E. coli* flagellar genes in nutrient-rich medium. **a**, Schematic representation of the *flhDC* operon in strain MG1655, with native (MG1655 *WT*) or inducible (*Ptac*) regulation of expression. The native regulatory region of the *flhDC* operon, including the upstream *IS1H* insertion element, was replaced in the *Ptac* strain with the *tac* promoter inducible by isopropyl β-d-1-thiogalactopyranoside (IPTG); an additional copy of the *lacI* gene (Lac repressor) was inserted upstream of the *tac* promoter to reduce the basal expression. **b,** Flow cytometry measurement of P*_fliC_*-GFP reporter activity in mid-exponential cultures of MG1655 *WT* or its *Ptac* derivative grown in tryptone broth (TB) medium. Flagellar gene expression in the *Ptac* strain was induced with the indicated concentrations of IPTG (in µM). Flow cytometry histograms of three biological replicates (*n =* 3) are shown as violin plots in different hues (AU – arbitrary units). **c,** P*_fliC_* reporter activity determined either as the median GFP intensity at mid-exponential growth phase in flow cytometry (FC) measurements (black symbols) or as the peak of GFP expression normalized by OD_600_ in plate reader (PR) cultures (red symbols). Both data sets were aligned by MG1655 *WT* expression (horizontal dashed line). Points are the mean values (*n* = 3) and error bars are the standard deviations (mean ± s.d.). **d**, Dependence of the average cell swimming velocity in cultures of the indicated *E. coli* strains on the activity of the P*_fliC_* reporter as determined by flow cytometry. The average swimming velocity was calculated as the product of the swimming fraction and the swimming velocity of motile cells (see Extended Data Fig. 1c,d for individual values). Motility and reporter expression were determined separately for each replicate culture (indicated by individual symbols). **e**, The growth fitness cost of flagellar gene expression. Fitness cost was determined as the percentage of cells (in %) of either the MG1655 *WT* or *Ptac* strain induced by different concentrations of IPTG in co-cultures with the non-flagellated *ΔflhC* strain after 24 h of growth with shaking (200 rpm) in TB medium. The strains were initially co-inoculated in a 1:1 ratio. P*_fliC_* activity measured in the plate reader was used to plot the data; mean ± s.d. (*n =* 3) is shown for both parameters.

To understand how motility changes as a function of gene expression, we characterized swimming behavior in populations of MG1655 *WT* and *Ptac* cells using differential dynamic microscopy^27^ (see Supplementary Note 1 and Extended Data Fig. 2). We observed that population-averaged cell swimming velocity initially increased with expression at low levels of induction, but saturated at high levels of expression (Fig. 1d). Notably, this saturation occurred around the level of motility gene expression seen in the wild-type strain. A similar pattern was observed when the fraction of well-swimming cells within the population, as determined by our motility assay, and the swimming velocity of only these cells were plotted individually (Extended Data Fig. 1c, d). The cell swimming velocity at the highest expression level was even slightly reduced (Fig. 1d and Extended Data Fig. 1c). Two other derivatives of *E. coli* K-12, W3110^28^ and RP437 (the latter is commonly studied as a wild type for *E. coli* chemotaxis^29^), both showed a similar relation between flagellar gene expression and motility, but were slightly less motile than MG1655 *WT* (Fig. 1d). The poorer swimming performance of RP437 may be a consequence of its extensive mutagenization^29^, and a previous study showed that the motility of this strain can be improved by experimental evolution^20^.

We further investigated the effect of motility on fitness by co-culturing CFP-labeled MG1655 *WT* or *Ptac* strains with a non-flagellated YFP-labeled *ΔflhC* strain. The fitness cost of flagellar regulon activity over a culture passage was determined as the reduction in relative cell number of the tested strain in the co-culture from the initial 50% at inoculation^20,21^. This cumulative fitness cost gradually increased with the level of motility genes expression over the entire range of induction tested (Fig. 1e). Thus, expression of motility genes beyond the native level in *E. coli* K-12 strains does not appear to provide any additional benefit, but nevertheless imposes an increasing fitness cost.

### Hydrodynamic constraints limit cell velocity at high levels of flagellar production

The saturation of *E. coli* motility at high levels of flagellar gene expression could be due either to some bottleneck in the biogenesis of functional flagella or to limits in the physical propulsion by multiple flagella. To distinguish between these two possibilities, we first determined how the activity of the flagellar regulon corresponds to changes in flagellation. Staining flagella with an amino-specific fluorescent dye^30^ revealed a clear dependence of the number and length of flagella on the expression of the flagellar regulon (Fig. 2a). The average number of flagellar filaments per cell showed an approximately linear increase with the activity of the P*_fliC_* reporter (Fig. 2b, Extended Data Fig. 3a). The length of flagellar filaments also showed a moderate increase followed by an apparent saturation (Fig. 2c, Extended Data Fig. 3b). These results were consistent with increased amounts of intra- and extracellular flagellin, determined by immunoblotting (Extended Data Fig. 4). Thus, *E. coli* cells can synthesize more flagella at levels of motility gene expression that exceed those of wild-type cells, but this increase does not translate into higher swimming velocity.

**Fig. 2.**
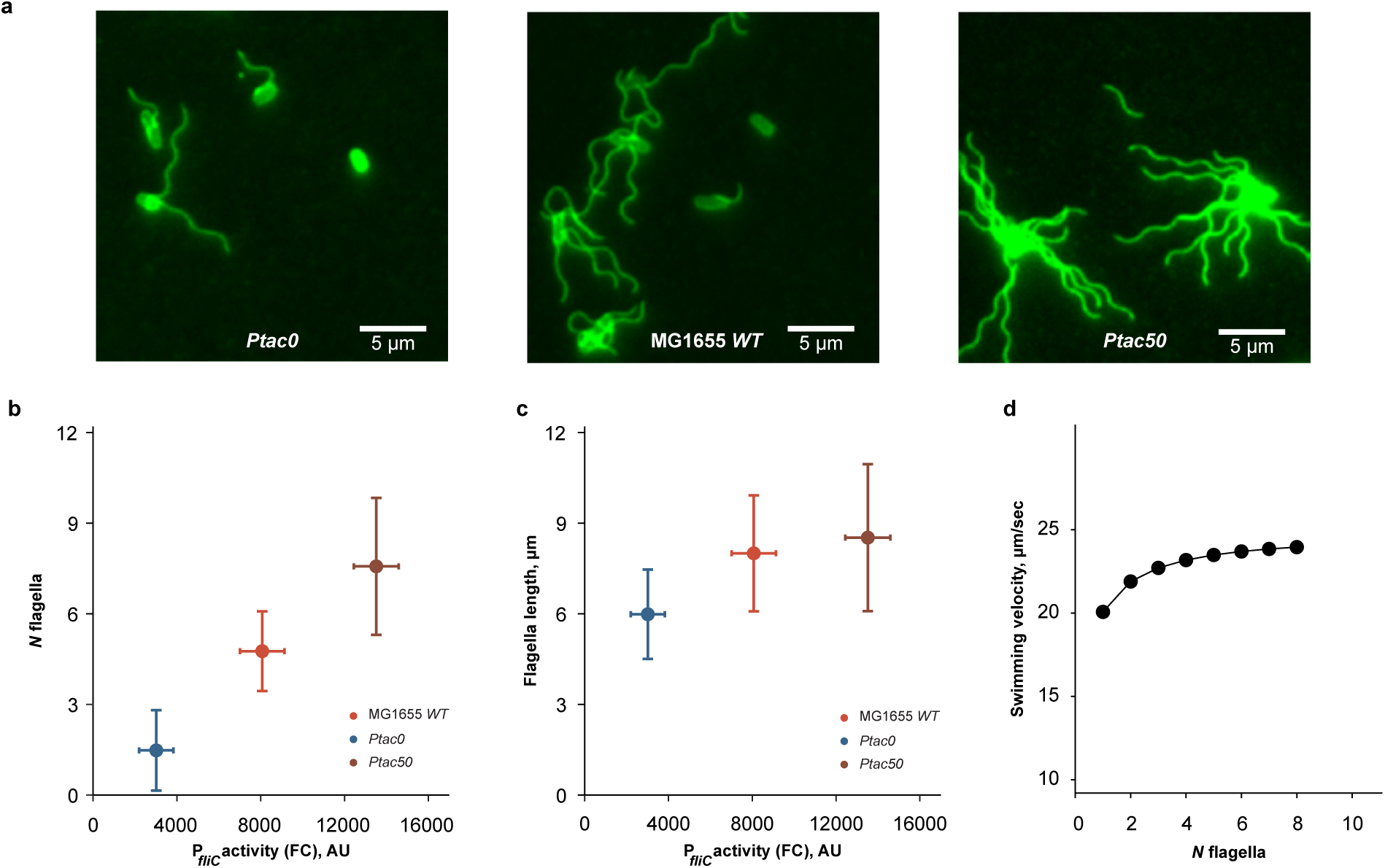
Limitation of *E. coli* motility at high expression of flagellar genes. **a-c**, Changes in *E. coli* flagellation with varying expression of flagellar genes. Fluorescence microscopy images of MG1655 *WT* or *Ptac* cells grown either without (*Ptac0*) or with 50 μM IPTG (*Ptac50*), stained with amino-specific fluorescent dye to visualize flagella (**a**). Corresponding quantification of the number (**b**, *N* flagella) and length (**c**, in µm) of flagella as a function of P*_fliC_* activity measured by flow cytometry (FC, *n* = 3 biological replicates, mean ± s.d.). Data from the same experiments were used to quantify both the number and length of flagella; *n* = 100 cells from different fields of view (**b**) and *n* = 35-50 flagellar filaments in 10-20 cells (**c**). See Extended Data Fig. 3 for value distributions and significance analysis. **d**, Dependence of swimming velocity on the number of flagellar filaments, predicted by the RFT physical model of the multi-flagellated microswimmer (see Supplementary Note 2 for details). Our RFT model takes into account that cells with a higher number of flagella also have longer filaments, as observed experimentally.

Alternatively, this saturation of swimming with flagella number could be explained by the physics of *E. coli* motility. The hydrodynamics of flagella-propelled bacterial swimming is well understood and can be captured by relatively simple mathematical models such as resistive force theory (RFT)^31,32^. We therefore used RFT to describe the swimming of a multi-flagellated bacterium, where multiple flagella form a tight bundle that rotates to propel the cell (Supplementary Note 2 and Extended Data Fig. 5a). Based on our experimental measurements (Extended Data Fig. 5b,c), we assume that the flagellar motors operate at a constant speed that does not depend on the number of flagella, which may be the maximum speed of the motor torque-speed relationship. Indeed, the load per motor is low and decreases as the number of flagella increases (Extended Data Fig. 5g), because now multiple motors share the torque generation necessary for bundle rotation and cell propulsion. Our model predicts that swimming velocity should initially increase with motility gene expression and then saturate, in agreement with the experimental data (Fig. 2d and Extended Data Fig. 5d-f). The initial increase stems from the increase of flagellar length and the increased thickness of the bundle formed by more flagella. Saturation then occurs in the RFT model at high number of filaments because the viscous drag of the cell body becomes negligible compared to the drag of the flagella themselves. As a consequence, any increase in thrust resulting from adding more flagella is offset by an equal increase in viscous drag, since the two have identical dependencies on flagellar length and bundle thickness. Although our model is clearly simplified, in particular, does not capture all the complexity of flagella bundle hydrodynamics^33^, it strongly indicates that the ability of *E. coli* to increase its swimming velocity by increasing the number and length of flagella is indeed limited by the hydrodynamics and mechanics of flagellar propulsion in viscous media.

### Motility gene expression follows the potential benefit of chemotaxis under carbon-limited conditions

Since expression of the flagellar regulon is under catabolite repression during carbon-limited growth in minimal media, we asked whether this regulation serves to maximize swimming, as observed in nutrient-rich medium, or whether it optimizes an alternative target. Consistent with its C-line-dependent regulation^7,21,25^, the expression of motility genes in the MG1655 *WT* strain grown in the minimal medium was much lower in the presence of a good (glucose) than a poor (succinate) carbon source (Fig. 3a). Expression in the *Ptac* strain at a given induction was also lower during growth on glucose, but this dependence was weaker, as expected for promoters that are not catabolite repressed^34^. Despite these differences, both swimming velocity (Fig. 3b) and growth fitness cost (Extended Data Fig. 6) in the *Ptac* strain showed the same dependence on motility gene expression for both carbon sources. MG1655 *WT* levels also fit to this curve, but unlike growth in nutrient-rich medium, the native activity of the flagellar regulon clearly does not maximize swimming velocity in this case.

**Fig. 3.**
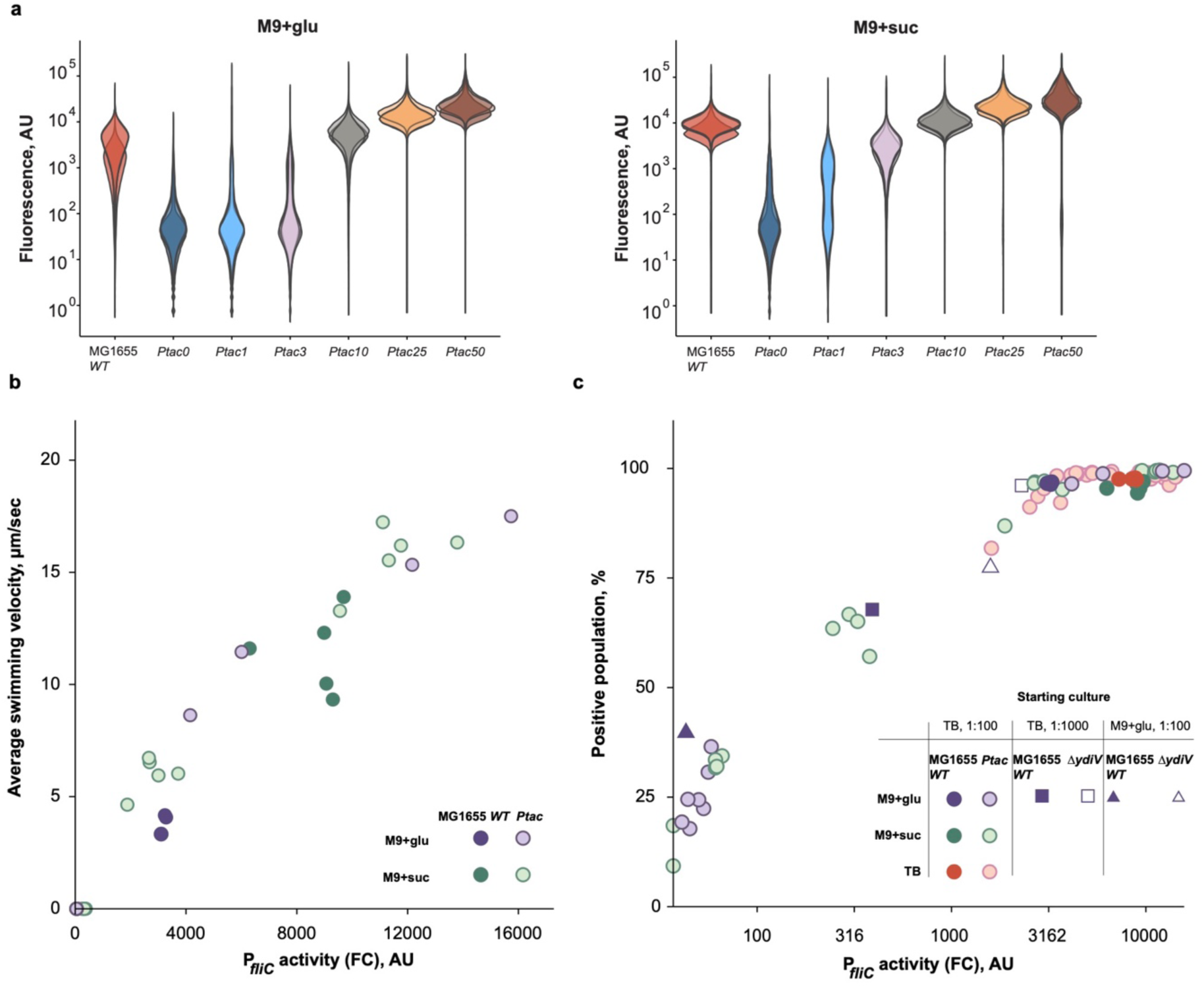
Motility of *E. coli* as a function of gene expression in minimal medium. **a**, Flow cytometry measurements of the P*_fliC_*-GFP reporter of MG1655 *WT* or *Ptac* strains grown to mid-exponential phase in M9 minimal medium, with either glucose (left) or succinate (right) as the sole carbon source. Labels are as in Fig. 1b. Flow cytometry histograms of three biological replicates are shown as violin plots in different hues (AU – arbitrary units). **b**, Dependence of average swimming velocity on the median P*_fliC_* reporter activity (flow cytometry, FC) for the indicated carbon sources and strains. Each dot represents an independent culture (biological replicate) for which both expression (P*_fliC_* reporter activity) and swimming were determined. **c**, Percentage of GFP-positive cells within the population of the MG1655 *WT*, *Ptac* and *ΔydiV* (lacks YdiV, the negative regulator of FlhDC; open symbols) strains as a function of median P*_fliC_* reporter activity, both measured by flow cytometry as in (**a**). Each symbol represents an independent culture. The inset describes different conditions used for the starting culture: the overnight culture pre-grown in TB (TB) or M9 glucose (M9+glu) was diluted to the fresh TB or M9 media (1:100 and 1:1000 indicate the dilution).

Instead, we hypothesized that native gene expression under carbon-limited growth might correlate with the potential benefit that could be achieved in a given carbon source by performing chemotaxis towards sources of additional nutrients, as proposed before^21^. Following this previous study, we measured the benefit of chemotaxis by providing localized sources of amino acids in co-culture between the *Ptac* strain (labeled with CFP) and its motile but non-chemotactic *ΔcheY* derivative (labeled with YFP) for different levels of motility gene induction (Extended Data Fig. 7). While the benefit of chemotaxis saturated at high levels of motility gene expression in both carbon sources, saturation occurred at much lower expression in the presence of glucose, with the point of saturation close to the native level of expression in the respective carbon source.

Another notable finding was the appearance of two distinct subpopulations, with almost negative and strongly positive expression, at low average levels of reporter activity in the *Ptac* strain (Fig. 3a and Extended Data Fig. 8). Interestingly, this separation appeared to be a function of the average reporter activity and did not depend on the carbon source (Fig. 3a and Fig. 3c). In this low expression range, the proportion of positive cells in the population increased up to a critical level of expression, after which the distribution became unimodal and it was rather the mean of the positive peak that increased with induction. Motility gene expression in MG1655 *WT* cells was above the critical level where bimodal behavior becomes apparent, even in culture grown on glucose. To investigate whether native regulation could also exhibit bimodality, we further reduced motility gene expression in wild-type cells by prolonged growth under catabolite repression in glucose, either by using a higher dilution of the TB-grown overnight culture or by pre-growing the overnight culture in glucose (Fig. 3c, Extended Data Fig. 9a). Indeed, both conditions reduced P*_fliC_* activity in the MG1655 *WT* cell population and revealed a bimodal pattern similar to that observed in the *Ptac* strain. Bimodality was also observed for a non-induced *Ptac* strain grown in TB (Fig. 1c and Fig. 3c). Thus, bimodality appears to depend solely on the expression level and not on the details of transcriptional regulation of the *flhDC* operon or on the growth medium.

Motility gene expression in *E. coli* has previously been shown to be pulsatile^26,35^ and this may be the cause of the observed bimodality. In the closely related species *Salmonella enterica,* motility genes are also known to exhibit bistable expression^36^. Both bistability (in *S. enterica*) and pulsatility (in *E. coli*) of expression were attributed to negative regulation of FlhDC activity by YdiV (RflP)^37^, with organism-specific differences in the topology of the YdiV regulatory circuit^35,37^. We therefore tested whether regulation by YdiV could be responsible for the emergence of bimodality in our experiments. As expected, the expression level of motility genes in a *ΔydiV* strain was elevated, and it was above the bimodality threshold in glucose even when the culture was inoculated from TB at a 1:1000 dilution (Fig. 3c and Extended Data Fig. 9b). However, when the expression level was sufficiently lowered by pre-growth in glucose, two distinct subpopulations could be clearly observed in the *ΔydiV* strain, suggesting that negative regulation by YdiV is not sufficient to explain the bimodal activation of the P*_fliC_* reporter.

### Activity of the flagellar regulon in natural isolates of *E. coli*

Finally, to investigate how investment in motility varies among *E. coli* strains that may have adapted to different ecological niches, we used the ECOR collection, which contains 72 isolates from different hosts and geographical regions^38^. From this collection, we first selected 61 strains that were sensitive to kanamycin and thus transformable with the P*_fliC_* reporter plasmid, and then discarded 23 non-swimming isolates that did not spread in porous (0.27%) TB agar. From the remaining 38 spreading isolates, a subset of 24 strains with moderate and good spreading abilities was chosen for further investigation (Supplementary Table 2).

Although the activity of the P*_fliC_* reporter varied widely among the TB-grown ECOR strains, it was consistently below or similar to that of the MG1655 *WT* strain (Fig. 4a and Extended Data Fig. 10a), indicating that the investment in motility by natural *E. coli* isolates is under similar limitation as in the K-12 strains. However, the swimming velocity of the majority of ECOR strains grown in liquid TB medium was lower than that of MG1655 *WT* and *Ptac* strains at similar levels of P*_fliC_* reporter activity (Fig. 4a and Extended Data Fig. 10a). Since previous studies showed that the motility of several pathogenic *E. coli* strains^39^ and other bacteria^40^ can be activated when cells are grown on a surface or in a porous medium, we measured the ability of ECOR strains to spread in porous 0.27% TB agar. Indeed, the spreading of most ECOR strains, including those that were poorly motile when grown in liquid, was comparable to that of MG1655 *WT* and *Ptac* (Fig. 4b).

**Fig. 4.**
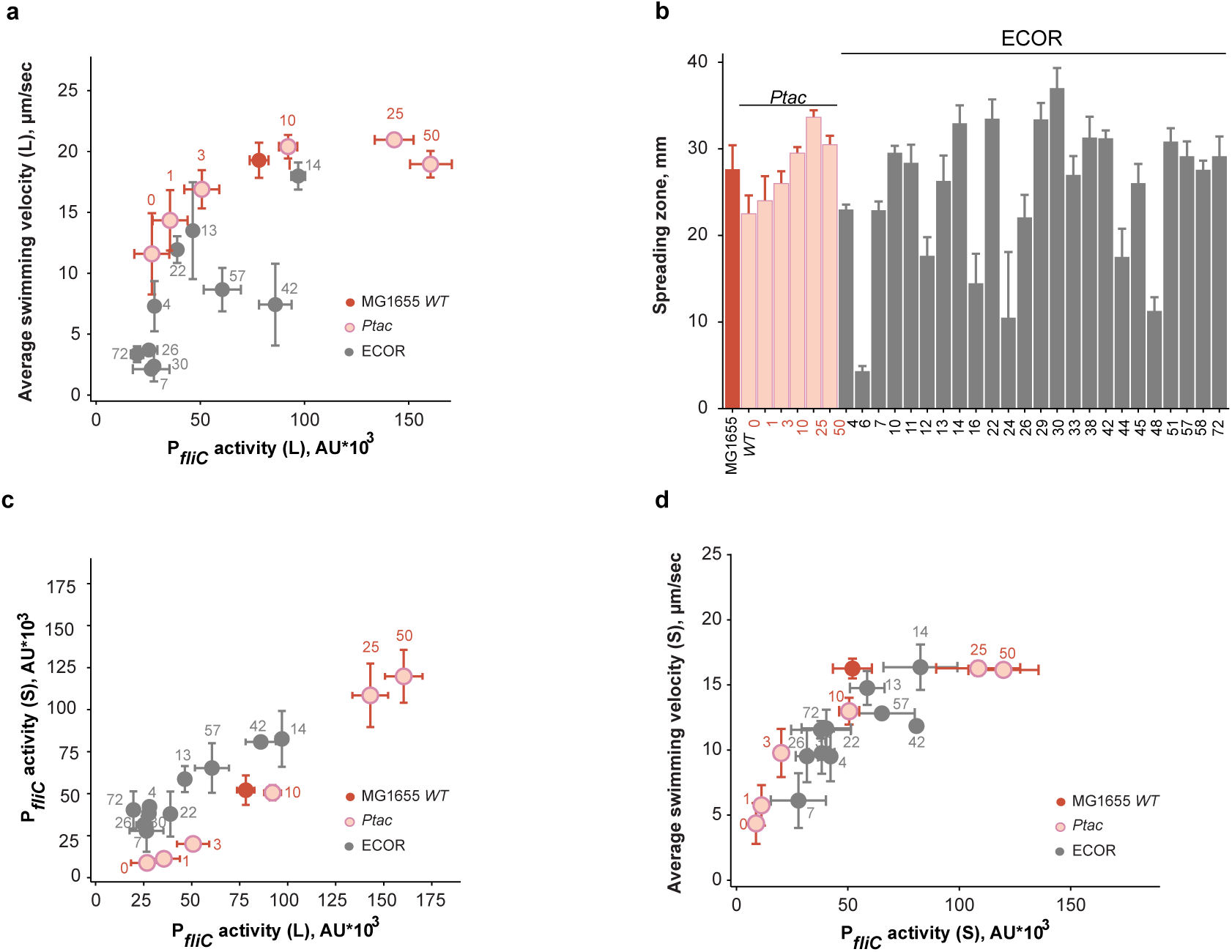
Motility of natural *E. coli* isolates. **a**, Relation between flagellar regulon activity and motility for representative ECOR strains (indicated here and throughout by their number in the collection) compared to MG1655 *WT* and *Ptac* strains; corresponding inducer concentrations (IPTG, µM) used for the *Ptac* strain are indicated by numbers in red. All *E. coli* cultures were grown in a liquid TB medium (indicated by L). The same mid-exponential cell culture was used to measure the P*_fliC_* reporter activity in the plate reader (GFP fluorescence normalized to OD_600_) and average swimming velocity (see Methods for details). Each point represents the mean value for both parameters (*n* = 3), with error bars indicating the standard deviations. **b**, Diameters of spreading zones formed by MG1655 *WT*, *Ptac* and ECOR strains in porous 0.27% TB agar, measured after 4-5 h incubation at 34°C (*n* = 3; mean ± s.d.). **c**, Correlation between P*_fliC_* reporter activity of in *E. coli* strains grown in liquid (L) or semi-solid (indicated by S) medium (0.5% TB agar) (*n* = 3; mean ± s.d.). **d**, Dependence of swimming velocity on P*_fliC_* activity for ECOR, MG1655 *WT* and *Ptac* strains grown on semi-solid (S) medium (*n* = 3, mean ± s.d.). Data for other ECOR strains are shown in Extended Data Fig. 10.

A possible explanation for this difference could be increased expression of motility genes in cells grown in porous media or on a semi-solid agar surface, where flagella rotate under high load^39,41–43^. We therefore measured the activity of the P*_fliC_* reporter in cultures grown on 0.5% TB agar plates. In this case, expression in individual strains correlated well with their spreading (Extended Data Fig. 10b). While we indeed observed an upregulation of reporter activity in such surface-grown compared to liquid-grown cultures for a few isolates (e.g. ECOR-72), this was not the case for the majority of ECOR strains (Fig. 4c, Extended Data Fig. 10c and Supplementary Table 2). However, when the motility of cells grown on an agar surface was subsequently analyzed in motility buffer (see Methods for details), the average cell swimming velocity was indeed higher for many ECOR strains compared to liquid-grown cultures, now showing a dependence of swimming velocity on expression similar to the MG1655 *WT* and *Ptac* strains (Fig. 4d, Extended Data Fig. 10d and Supplementary Table 2). Thus, the observed poor motility of many ECOR isolates grown in liquid medium cannot be generally explained by low activity of the flagellar regulon but rather indicates some deficiency in flagellar assembly or function in liquid-grown cell. Notably, however, both motility gene expression and swimming of all ECOR strains were always below or comparable to that of MG1655 *WT*, further supporting the fundamental nature of limitation imposed on *E. coli* motility by hydrodynamics.

## Discussion

How microorganisms regulate the allocation of their limited cellular resources under varying environmental conditions remains an open question. Although optimality theory^50^ predicts that gene expression levels should have been evolutionarily tuned to maximize an organism’s fitness, such optimization is a multifactorial problem with mostly uncharacterized constraints and trade-offs between conflicting optimization goals. Particularly challenging to understand are microbial strategies for allocating resources to costly functions that do not directly benefit growth or are not used under certain conditions, which can account for up to half of cellular protein resources^13,44,45^.

Here, we investigated resource allocation to flagellar motility, the most prominent of such non-growth related cellular functions in bacteria, by titrating the expression of the flagellar gene regulon and quantifying its impact on *E. coli* motility. We observed that the biogenesis of the motility apparatus, i.e., the number of flagella and their length, shows a dependence on gene expression over a wide range, demonstrating that *E. coli* can increase its flagellation beyond the level observed in wild-type strains with the native regulation of gene expression. The effect on growth fitness increases proportionally with resource investment, too, consistent with flagella biosynthesis being the major component of motility costs^20,21^. In contrast, cell swimming velocity increases as a function of motility gene expression until the number of flagella reaches ∼5, but saturates above this level. This dependence of swimming velocity on the number and length of filaments was well captured by a mathematical model describing the swimming of a multi-flagellated bacterium using the resistive force theory, suggesting that the observed saturation of cell velocity is the consequence of hydrodynamic constraints on *E. coli* motility. Further supporting the general nature of this relation, not only the K-12 strains, but also the majority of motile natural isolates of *E. coli* mapped to the same unique expression-swimming relation under conditions that favored their motility.

Strikingly, although the activity of the flagellar regulon differed among the wild-type *E. coli* strains tested and between conditions, it was invariably confined to the sub-saturating part of the expression-swimming relation. In a fraction of the strains, including K-12 derivatives and several natural isolates, motility gene expression in the nutrient-rich medium was most likely selected to maximize swimming velocity. This could indicate a high importance of swimming, e.g., for colonization of the environment^19,46^. However, even in these strains, expression levels remain bounded by the critical level at which swimming velocity saturates, indicating that cells avoid unnecessary resource expenditures that provide no additional benefit. Expression levels in other *E. coli* isolates map to different points on the expression-swimming curve, covering the range below saturation of motility. Such heterogeneity could be due to different selection pressures on motility in the ecological niches occupied by these isolates, which is consistent with findings that differences in motility allow coexistence and niche segregation between *E. coli* strains, both *in vitro*^25^ and in an animal host^47^.

While many *E. coli* strains, including the K-12 derivatives and some natural isolates, swim similarly well when grown in either liquid or porous media, we observed that most natural isolates showed good motility only when grown in porous or semi-solid media, possibly reflecting conditions in the animal gut. The mechanism underlying this effect needs to be further characterized, but it does not seem to be explained by a previously reported mechanosensing-based upregulation of the entire flagellar gene regulon in porous media^39^. Many *E. coli* isolates swim poorly when grown in liquid despite having comparatively high activity of the flagellar regulon, and only achieve the motility expected based on their gene expression when grown on semi-solid medium. For these isolates, growth in liquid may result in the assembly of poorly functional motors or flagella. A potential mechanism for such flagellar motor remodeling in *E. coli* could be the previously described recruitment of additional force-generating units under load^41,43^, but it remains to be seen whether this recruitment is sufficiently long-lasting to account for these isolates retaining high motility even after transfer to a liquid environment.

When grown under carbon limitation, *E. coli* cells exhibited similar expression-swimming and expression-cost relations in both good and poor carbon sources, despite expected growth-dependent changes in cell size^23^. However, under these conditions, native expression of *E. coli* motility genes clearly does not maximize swimming. Instead, it correlates well with saturation of the benefit that *E. coli* could derive from chemotaxis-dependent accumulation to sources of additional nutrients, consistent with the strategy of anticipatory investment in motility^21^.

The reduced activity of the flagellar regulon under carbon-limited growth revealed another prominent feature of its regulation in *E. coli*, namely the appearance of two distinct subpopulations of cells below a certain threshold of average P*_fliC_* reporter activity. This bimodality may be related to the recently described pulsatile activation of flagellar genes in *E. coli* at intermediate expression levels of the master regulator FlhDC^26,35^. However, whereas this previous work concluded that pulsatility of expression is caused by the negative regulation of FlhDC by YdiV^26^, this regulation was not sufficient to explain the bimodality in our experiments. Furthermore, based on the established quantitative relation between gene expression and swimming motility, we could speculate on possible physiological reasons for such differentiation into distinct subpopulations. The bimodality of gene expression in microorganisms is commonly interpreted as stochastic bet-hedging behavior, which may be a better strategy in an unpredictable environment than a single adaptive phenotype^48–50^. While similar arguments were used to rationalize the differentiation of a bacterial population into motile and non-motile phenotypes^26,35,36^, here we propose a different, though not mutually exclusive, explanation. We noticed that the bimodality in our experiments occurs at the average expression that is below the level that would correspond to approximately two flagella per cell. Given that swimming with fewer than two flagella becomes inefficient, we argue that the observed bifurcation serves to avoid this “average”, poorly motile phenotype, which is unable to benefit from motility but still pays the fitness cost. Such “enforced” bet hedging may provide an alternative explanation for evolutionarily selected bimodality of gene expression, which is likely to apply not only to bacterial motility, but also to other cases where an intermediate phenotype is less fit than either of the extreme phenotypes. Thus, the hydrodynamics of flagella-mediated motility may not only determine the upper limit of swimming velocity at high levels of motility gene expression, but may also explain its bimodality at low levels of expression.

## Methods

### Strains and growth conditions

All *E. coli* strains, including natural isolates from the *E. coli* Reference Collection (ECOR)^38^ and plasmids used in this study are described in Supplementary Tables 1 and 2. The strain with inducer-dependent expression of *flhDC* operon (*Ptac*) was constructed previously^21^ by replacing the native regulatory region of the *flhDC* operon, including the upstream *IS1H* insertion element, in the MG1655*Δflu* background with the *tac* promoter inducible by isopropyl β-d-1-thiogalactopyranoside (IPTG). To reduce the basal expression of the *flhDC* operon, the *lacI* gene encoding the Lac repressor was additionally inserted upstream of the *tac* promoter. Deletion of the *ydiV* gene in MG1655*Δflu* and its *Ptac* derivative was performed by P1 transduction from the KEIO collection^51^ followed by curation of the resistance cassette by FLP recombination^52^. Deletion of the *flu* gene encoding the major *E. coli* adhesin, antigen 43, in the MG1655 group strains was used to prevent autoaggregation of motile planktonic cells^53^ and thus facilitate subsequent characterization of motility^21^.

To evaluate the activity of the flagellar regulon, strains were transformed with the plasmid carrying the GFP reporter for *fliC* promoter (P*_fliC_*) as described previously^21^. For pairwise growth competition experiments, performed as before^21^, the strains were labeled by expression of either cyan or yellow fluorescent proteins (CFP or YFP) from the pTrc99a vector under the control of the IPTG-inducible synthetic P*_trc_* promoter^54^. Since pTrc99a carries an extra copy of *lacI*, which reduces the leaky expression from the genomic P*_tac_* promoter and thus the inducibility of expression in the *Ptac* strain, an empty pTrc99a vector was transformed into *Ptac* and other *E. coli* K-12 strains for comparability.

*E. coli* strains were grown in either lysogeny broth (LB; 10 g l^−1^ of tryptone, 5 g l^−1^ of yeast extract, 5 g l^−1^ of NaCl), tryptone broth (TB; 10 g l^−1^ of tryptone, 5 g l^−1^ of NaCl), and either M9 (5× stock made with 64 g l^−1^ of Na_2_HPO_4_-7H_2_O, 15 g l^−1^ of KH_2_PO_4_, 2.5 g l^−1^ of NaCl, 5.0 g l^−1^ of NH_4_Cl, 2 mM MgSO_4_, 0.1 mM CaCl_2_, 1μM FeSO_4_, and 1μM ZnCl_2_) or Tanaka (34 mM Na_2_HPO_4_, 0.3 mM MgSO_4_, 64 mM KH_2_PO_4_, 10 μM CaCl_2_, 1μM FeSO_4_, and 1μM ZnCl_2_)^55^ minimal media supplemented with 0.4% glucose or 15 mM succinate as the sole carbon source. Ampicillin (100 μg ml^-1^) and/or kanamycin (100 μg ml^-1^), and isopropyl β-d-1 thiogalactopyranoside (IPTG) were added to the media when necessary.

### Reporter activity measurements

P*_fliC_* reporter activity was measured by either flow cytometry or plate reader assay. Unless otherwise stated, for flow cytometry, overnight cultures grown in TB (37°C, 200 rpm) were diluted 1:100 in 10 ml of the respective target medium. When minimal medium was used, cells were washed three times in medium without carbon source before inoculation. Cultures were incubated at 34°C with shaking (270 rpm) and harvested at mid-exponential phase (OD_600_ = 0.4-0.6 for TB or 0.3-0.5 for M9). Cultures were diluted ∼50-fold in tethering buffer (6.15 mM K_2_HPO_4_, 3.85 mM KH_2_PO_4_, 0.1 mM EDTA, 1 µM methionine, 10 mM sodium lactate, pH 7.0) and fluorescence was detected using a 488 nm laser (100 mW) and a 510/20 nm bandpass filter for GFP on a BD LSRFortessa SORP cell analyzer (BD Biosciences, Germany). 30,000 individual events were analyzed in each experimental run. Gating was first performed on an FSC-A/SSC-A plot and on an SSC-W over SSC-H plot to exclude doublets. Events in the samples with fluorescence intensities higher than the background signal from the MG1655 *WT* or *Ptac* strain without the reporter plasmid were considered ‘positive’. The proportion of ‘positive’ events per sample and summary statistics (mean, median fluorescence values) of both the ‘positive’ and the ‘whole’ population were assessed during the measurements using BD FACSDiva^TM^ Software v8.0.1 during measurements. Data were collected in FCS 3.0 file format and analyzed using the flowCore package in R v. 4.2.2.

For growth and expression measurements in the BioTek Synergy H1 plate reader, cultures were inoculated into the 96-well plates (Greiner Bio-One) at a dilution of 1:1000 and grown at 34°C with double orbital shaking at a frequency of 548 cycles per minute (CPM) and a shaking amplitude of 2mm for 24 h (TB) or for 48-64 h (M9). GFP fluorescence was quantified using a monochromator-based filter set (excitation 485 nm, emission 530 nm, with a bandpass ≤18 nm for detection). Fluorescence and optical density (OD_600_) were measured every 10 min. For experiments shown in Extended Data Fig. 7, the TECAN Infinite M1000 PRO plate reader was used instead for consistency with the previous study^21^.

Reporter activity in ECOR isolates was measured after growth in liquid TB medium or on the surface of semi-solid TB agar (0.5%). For the liquid medium setup, day cultures were prepared in the same manner as for flow cytometry. For the semi-solid condition, 20 µL of the same overnight culture was spread on the surface of TB agar using glass beads. After drying for 15-20 min, the plates were incubated at 34°C for the same time as the strain grew in liquid medium until OD_600_= 0.4-0.6 (i.e., 2.5-4h). Cells were gently washed from the plates with 2 ml of motility buffer (6.15 mM K_2_HPO_4_, 3.85 mM KH_2_PO_4_, 0.1 mM EDTA, 67 mM NaCl, pH 7.0) and adjusted if necessary to final OD_600_ = 0.5, and 1 ml of a liquid-grown culture was also washed once in motility buffer. After another washing step, the cells were resuspended in 1 ml motility buffer supplemented with 1% glucose and 0.001% Tween-80. GFP fluorescence was measured in a TECAN Infinite 200 PRO plate reader at 480 nm wavelength, 9 nm bandwidth for excitation and 510 nm wavelength, 20 nm bandwidth for emission.

### Analysis of swimming velocity and flagella rotation

Bacterial cell motility was analyzed as previously described^21,56^. Briefly, 1 ml of the same cell culture as prepared for flow cytometry was gently centrifuged (4000 rpm, 5 min), washed twice in motility buffer, and resuspended in 1 ml motility buffer supplemented with 1% glucose and 0.001% Tween-80. 3-5 µL of this cell suspension was introduced into a custom-made chamber between two coverslips, and motility was imaged by phase-contrast video-microscopy (Nikon TI Eclipse, 10x objective with NA = 0.3, Phase 1 ring, CMOS camera EoSens 4CXP), with 10,000 frames being recorded at a rate of 100 frames per second (fps). Motility parameters, in particular the fraction of swimming cells and the swimming velocity of the swimmers, are extracted from the movies using differential dynamic microscopy (DDM)^55^ (see Supplementary Note 1).

To determine the frequency of flagella rotation, samples were prepared in the same manner as described for swimming velocity analysis. A 10,000-frame movie with a field of view of 512 x 512 px^2^ (1 px = 0.7 µm) was acquired far from the sample surfaces under dark field illumination (Nikon TI Eclipse, 10x objective with NA = 0.3, CMOS camera EoSens 4CXP) at a rate of 800 fps. Dark field illumination is obtained by combining an aligned Ph3 condenser ring with the 10x objective on the Nikon TI Eclipse microscope. All data were analyzed using the dark field flicker microscopy (DFFM) method^57^ (see Supplementary Note 1) implemented in ImageJ (https://imagej.nih.gov/ij/) with custom-written plugins. Briefly, DFFM uses the flickering that results from changes in the direction in which light is scattered by anisotropic objects as they rotate to measure the rotation speeds of the cell body and flagella.

### Motility assay in soft agar

Motility driven spreading of *E. coli* in 0.27% TB soft agar was analyzed as previously described^39^. Briefly, 2 µl of overnight cultures grown in TB (37°C, 200 rpm) were transferred to the soft agar plates, and the diameters of the spreading zones were measured after 4-5 h of incubation at 34°C by capturing images with an iPad camera and quantifying the diameter of the spreading zone using ImageJ.

### Pairwise growth competition

Growth competition assays were performed as previously described^21^. Briefly, the overnight cultures of the MG1655 *WT* or *Ptac* strain expressing CFP and the *ΔflhC* strain expressing YFP, grown individually in TB (37°C, 200 rpm), were mixed in a 1:1 ratio to final OD_600_ = 0.0025 in 2.5 mL of fresh media and cultured for 24 h (TB) or 48-72 h (M9 minimal medium) at 34°C and 200 rpm. The expression of YFP and CFP was induced with 10 µM IPTG for the co-culture containing the MG1655 *WT* strain or by the corresponding IPTG concentrations used for induction of the chromosomal *Ptac* promoter. For the chemotactic benefit assay, differentially labeled non-chemotactic *ΔcheY* strain and MG1655 *WT* or *Ptac* strains were grown in Tanaka minimal medium for 72 h without shaking in the presence of nutrient gradients generated by 40 µLlarge agarose beads (2% agarose) containing 12% casein hydrolysate as described previously^21^. The initial and final proportions of CFP- and YFP-labeled cells were measured by flow cytometry on the BD LSRFortessa SORP cell analyzer (BD Biosciences). The sample was excited with lasers at 447 nm (75 mW), 514 nm (100 mW), and 488 nm (20 mW), with the latter used to identify all cells. CFP and YFP emission signals were detected at 470/15 nm and 542/27 nm, respectively. The fraction of CFP/YFP-‘positive’ events per sample was assessed during the measurements using BD FACSDiva^TM^ Software v8.0.1. Summary statistics were collected in csv file format and analyzed in R v. 4.2.2.

### Measurements of flagellar length and number

For flagella staining, 1 ml of the mid-exponential cell culture grown in TB as described above was centrifuged (3000g, 3 min) and gently washed three times in Buffer A (10 mM KPO_4_ buffer, 0.1 mM EDTA dipotassium salt, 67 mM NaCl, 0.001% Tween-80, pH 7.0). The cell pellet was resuspended in 400 µL of Buffer B (same as Buffer A but adjusted to pH 7.8 with NaHCO_3_), and 8 µl of 10 µg ml^-1^ Alexa Fluor 594 succinimidyl ester dye dissolved in DMSO was added to the mixture. Samples were incubated at 30°C in the dark with gentle shaking (100 rpm) for 90 min, washed three times in Buffer A and diluted fivefold in Buffer A. 3-5 µl of cell suspension was applied to a 1% agarose pad (in tethering buffer) and transferred to a 2-well µ-Slide (ibidi, Germany).

Fluorescence widefield images were acquired using a Zeiss Elyra 7 inverted microscope with a 63x oil/1.46 oil objective and a further 1.6X magnification. The sample was excited with a 561 nm 500 mW laser (1% power) using a quadruple band dichroic and emission filter. The fluorescence emission of the succinimidyl ester was detected at 595/50 nm interval with a PCO 4.2 Edge sCMOS camera, the exposure time was 100 ms. The number of flagella was quantified for randomly selected 100 cells in multiple fields of view, including both flagellated and non-flagellated cells. The length of flagellar filaments (35-50 filaments per condition) was measured using segmented line tool of ImageJ.

### Immunoblot analysis of intra- and extracellular flagellin

To shear flagellar filaments, a 1 ml aliquot of the mid-exponential cell culture was passed through a 1 ml syringe with the 26G needle 20 times, and centrifuged at 2500 g for 10 min. The supernatant and cell pellet, resuspended in 333 µL of TB medium, were further analyzed by immunoblot. To transfer the samples to the membrane after SDS-PAGE, a PerfectBlue Semi-Dry Electroblotter (Peqlab, VWR, Germany) was used at constant amperage for 1 h (150 mA for 8*6 cm membrane and 1.5 mm thick gel). After transfer, the membrane was stained with Revert™ 700 Total Protein Stain for Western Blot Normalization (LI-COR Biosciences, Germany) and, after blocking,incubated overnight (4°C, orbital shaking) with the primary anti-flagellin antibody (Antikoerper, Germany) diluted 1:10000 followed by the secondary IRDye 800CW anti-rabbit IgG antibody (LI-COR Biosciences, Germany) antibody at a dilution of 1:10000. Fluorescence was measured using an Odyssey Clx Infrared Imaging System (LI-COR Biosciences, Germany) in two channels (700 and 800 nm). Images were analyzed and processed using ImageJ.

### The model of flagellum-mediated bacterial swimming

The model for multiflagellated propulsion extends the classical force balance analysis for uniflagellated propulsion^31,58^ and accounts for our measurements of swimming speed, cell body rotation speed, and flagellar rotation speed, as well as flagellar length, flagellar number, and cell size. The model is described in detail in Supplementary Note 2. Briefly, we assume that the *N* flagella form a single tight bundle, described in the framework of resistive force theory^31,59–61^ as a helix of larger thickness for a higher number of flagella, which is justified considering several macroscopic experiments at low Reynolds number with multiple helices^62,63^. We account for the increase in both flagellar length and flagellar number with increasing *flhDC* induction. The cell body is described as a counter-rotating rod^64,65^ of fixed size, consistently with our observation. The flagellar motor speed is assumed to be constant, in agreement with our measurements of the flagella and cell body rotation speeds. The balance of forces and torques acting on the cell body and the flagellar bundle provides predictions of the swimming speed and the rotation frequencies.

## Data and materials availability

All data are available in the main text or in Extended Data. All materials are available from the corresponding author upon request.

## Acknowledgments

We thank Julian Pietsch for careful reading of the manuscript. We thank Julian Pietsch and Santiago Kuhl for fruitful discussions. We thank Silvia Gonzalez Sierra and Gabriele Malengo for the technical assistance with flow cytometry and microscopy experiments, and Irina Kalita for the help with flagella labeling. This research was funded by the Max-Planck-Gesellschaft and by the Max Planck School Matter to Life supported by the German Federal Ministry of Education and Research (BMBF).

## Author contributions

I.L., B.N. and V.S. designed the study. I.L., R.C., B.N. and V.S. designed the experiments. I.L., R.C., H.Y., and B.N. performed the experiments. I.L., R.C, and H.Y. analysed the data. I.L., R.C., and V.S. wrote the manuscript.

## Materials & Correspondence

Correspondence and requests for materials should be addressed to Victor Sourjik.

---

**Extended Data Fig. 1.**
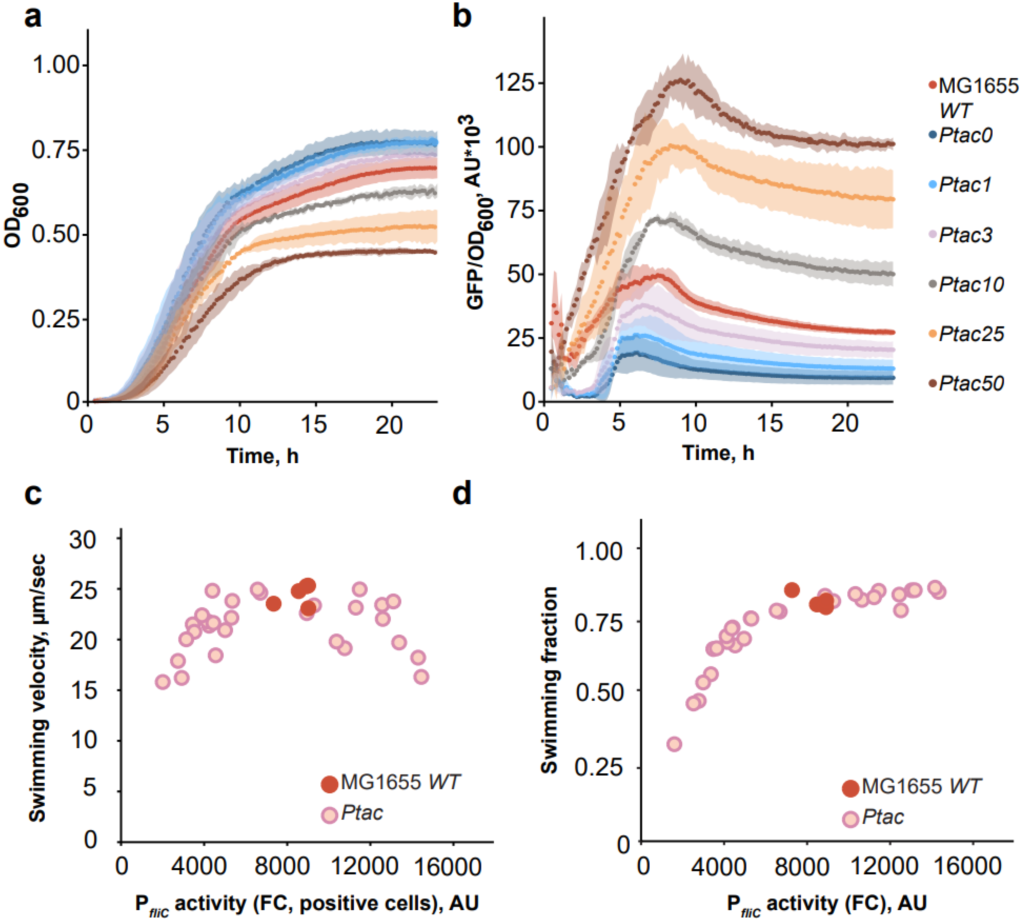
The effect of flagellar gene expression on growth and motility of MG1655 *WT* and *Ptac* strains in nutrient-rich medium (TB). Cell growth (a) and P*_fliC_* reporter activity (GFP/OD_600_) (b) were monitored in the indicated cultures for 24 h by measuring absorbance (OD_600_) and GFP fluorescence every 10 min in the plate reader. Numbers represent the corresponding IPTG concentration for the *Ptac* strain. Standard deviation is shown by the shaded area around the curves (*n* = 3 biological replicates, mean ± s.d.). Changes in the swimming velocity of motile cells (c) and swimming fraction (d) as a function of reporter activity measured by flow cytometry (FC). P*_fliC_* activity was determined as median GFP intensity only in GFP-positive cells (see Methods for details) (c) or in the whole population (d). Each point represents a single replicate culture.

**Extended Data Fig. 2.**
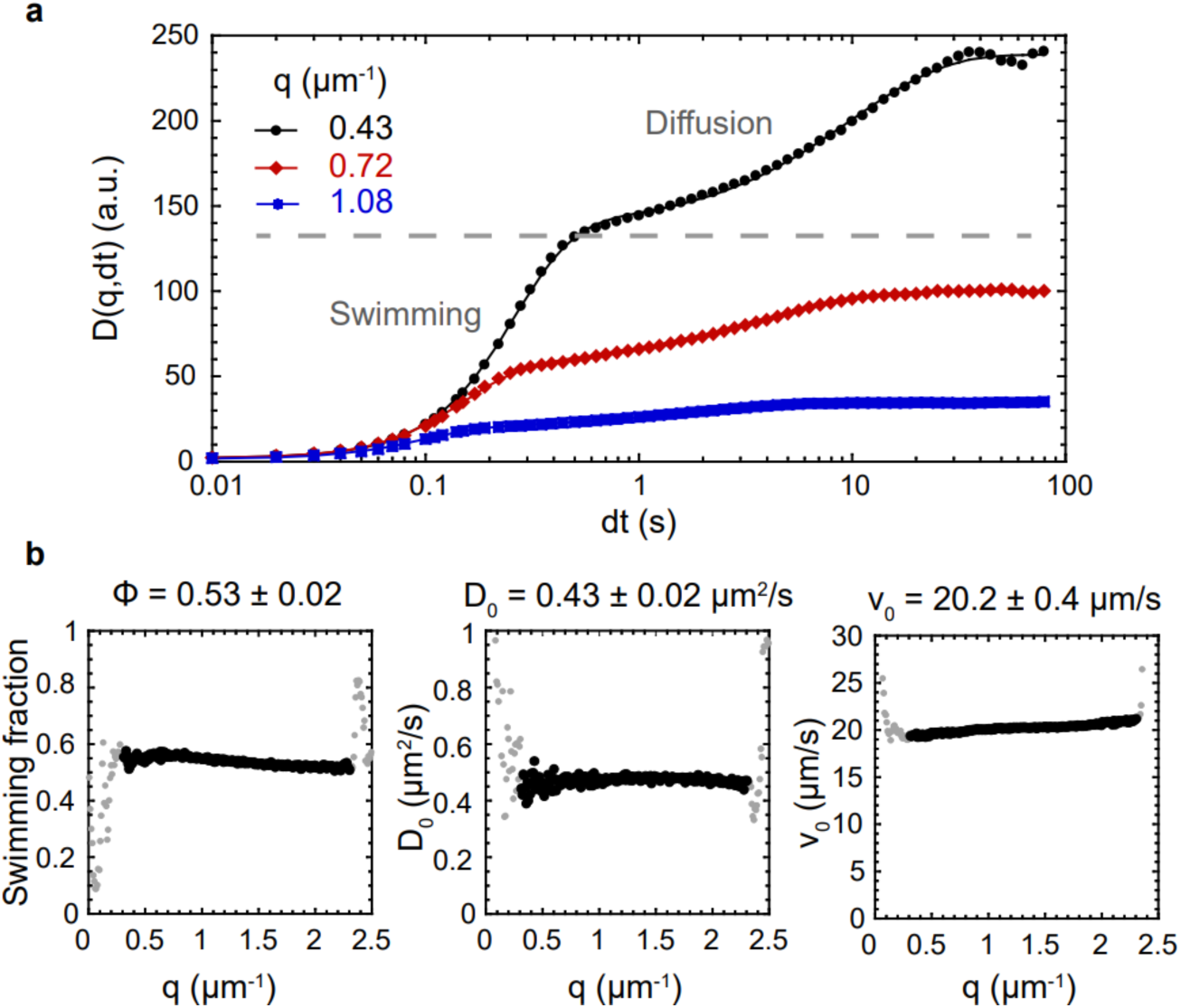
Differential dynamics microscopy analysis of cell motility. Shown is an example measurement for the *Ptac* strain at 0 µM IPTG induction. **a**, Differential intensity correlation functions (DICF) as a function of the lag time dt, for different values of the wave number q. The dashed gray line indicates the separation between the contribution of swimming (short times) and diffusion (long times) to the increase of the DICF. Points are experimental data and lines are fits by the (swimming + diffusion of non-swimmers) model (see Supplementary Note 1). **b**, Resulting fit parameters (fraction of swimmers, diffusion coefficient, and average velocity) as a function of q. Dark dots indicate successful fits and gray dots are the ones that fail due to either lack of full decorrelation (small q) or low signal over noise (large q). Consistent fit parameter values over the valid range of *q* validate the model. The mean and standard deviation of the fit parameter values over the valid range are indicated.

**Extended Data Fig. 3.**
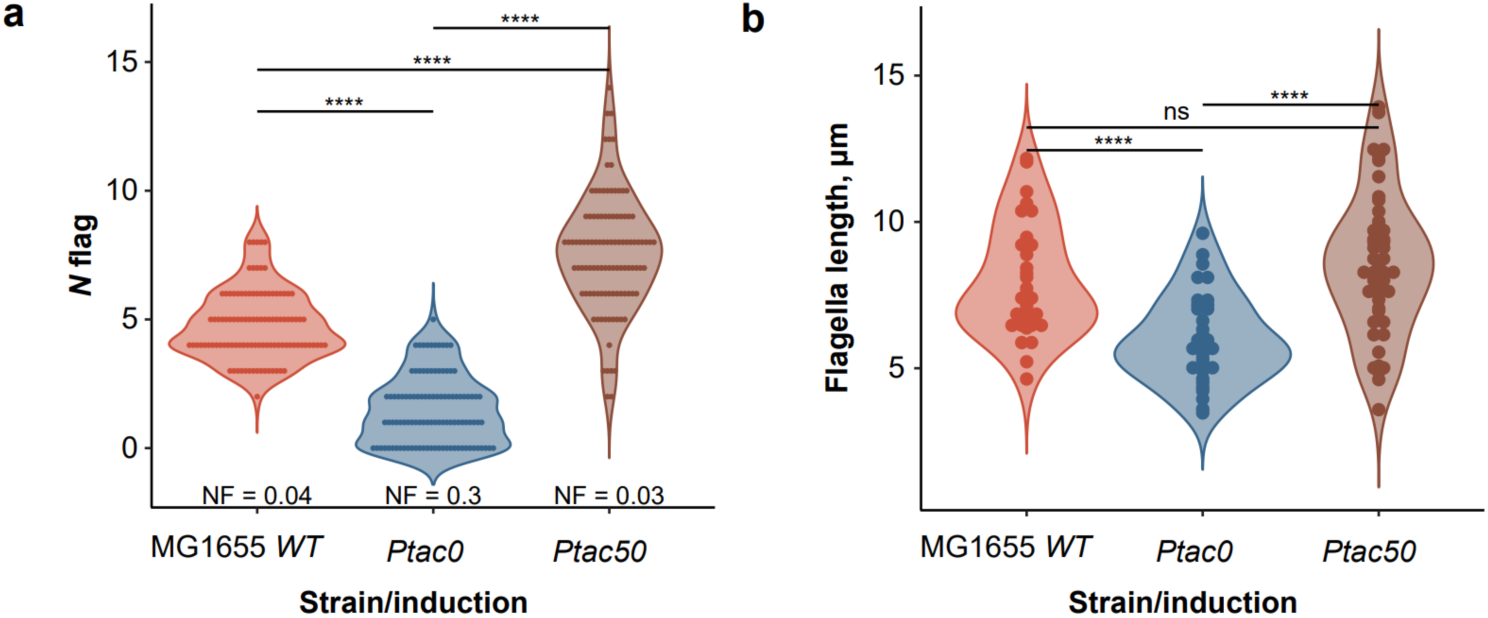
Distributions of flagella number and length in the population of MG1655 *WT* and *Ptac* strains (0, 50 μM IPTG). Each point on the violin plot is a single-cell measurement of flagella number (**a**, *n* = 100 randomly selected cells) or length (**b**, *n* = 35-50 flagellar filaments in *n =* 10-20 cells) for the indicated condition. Normality of means was tested by the Shapiro-Wilk test (*P* ≤ 0.05 (**a**), and *P* ≥ 0.05 (**b**)). Due to the large sample size (*n* > 20), a two-sided t-test was used for both (**a**) and (**b**) to compare the differences between the population means. Since the hypothesis of equal variances was rejected (Levene’s test, *P* ≤ 0.05), we used Welch’s t-test followed by the Holm-Bonferroni method to correct for multiple testing, and the adjusted *P* values (\*\*\*\**P* ≤ 0.0001) are shown on both panels. NF on (**a**) indicates the fraction of non-flagellated cells for each condition.

**Extended Data Fig. 4.**
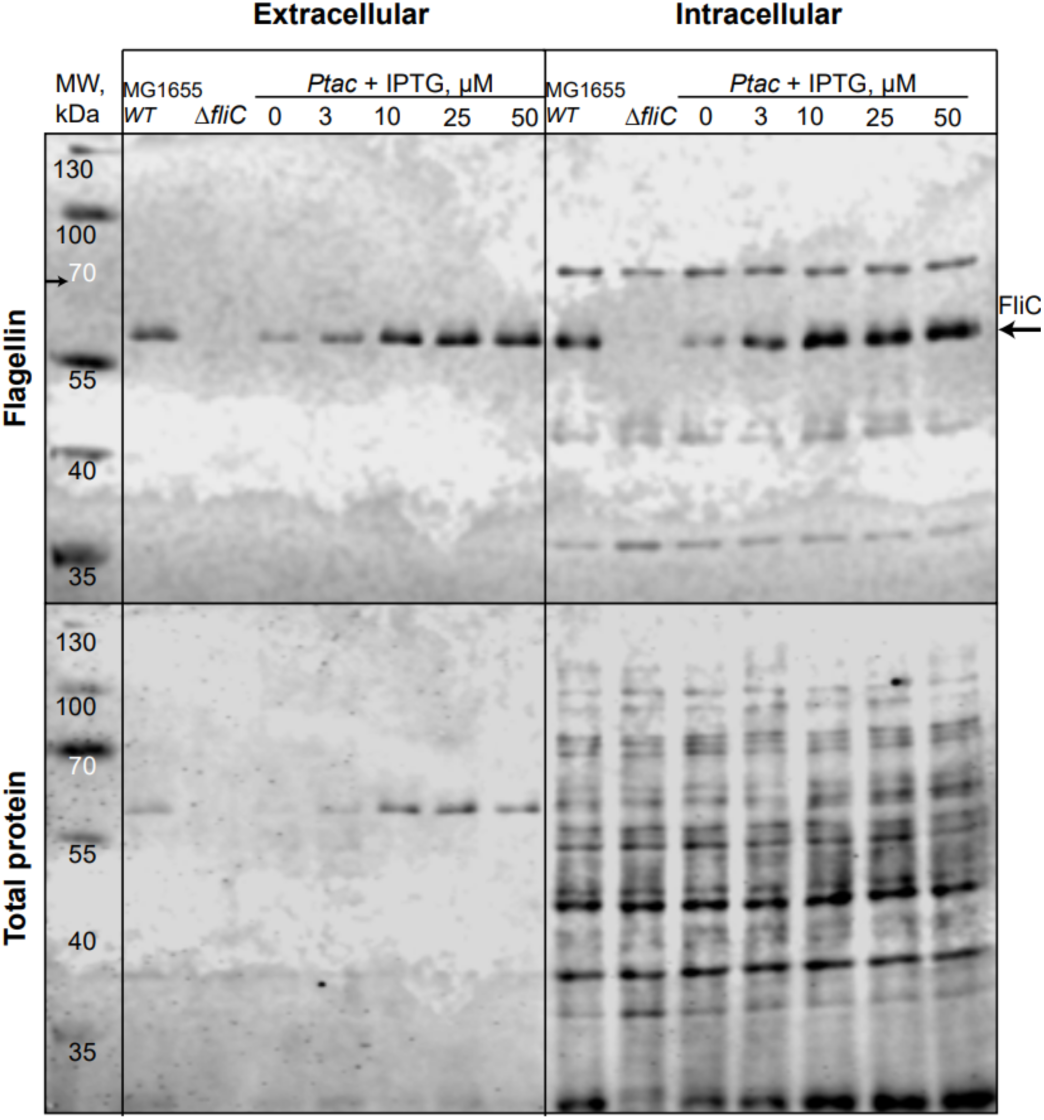
The amount of intra- and extracellular flagellin increases as a function of motility gene expression. Immunoblotting analysis of flagellin (FliC, indicated by black arrow) in intra- and extracellular fractions of MG1655 *WT*, *Ptac* and *ΔfliC* (negative control) cells. Sample volumes were adjusted by OD_600_ normalization prior to loading. Membrane staining for total protein was used as a loading control (bottom). MW, kDa – band profile of the prestained protein ladder. See Methods for details.

**Extended Data Fig. 5.**
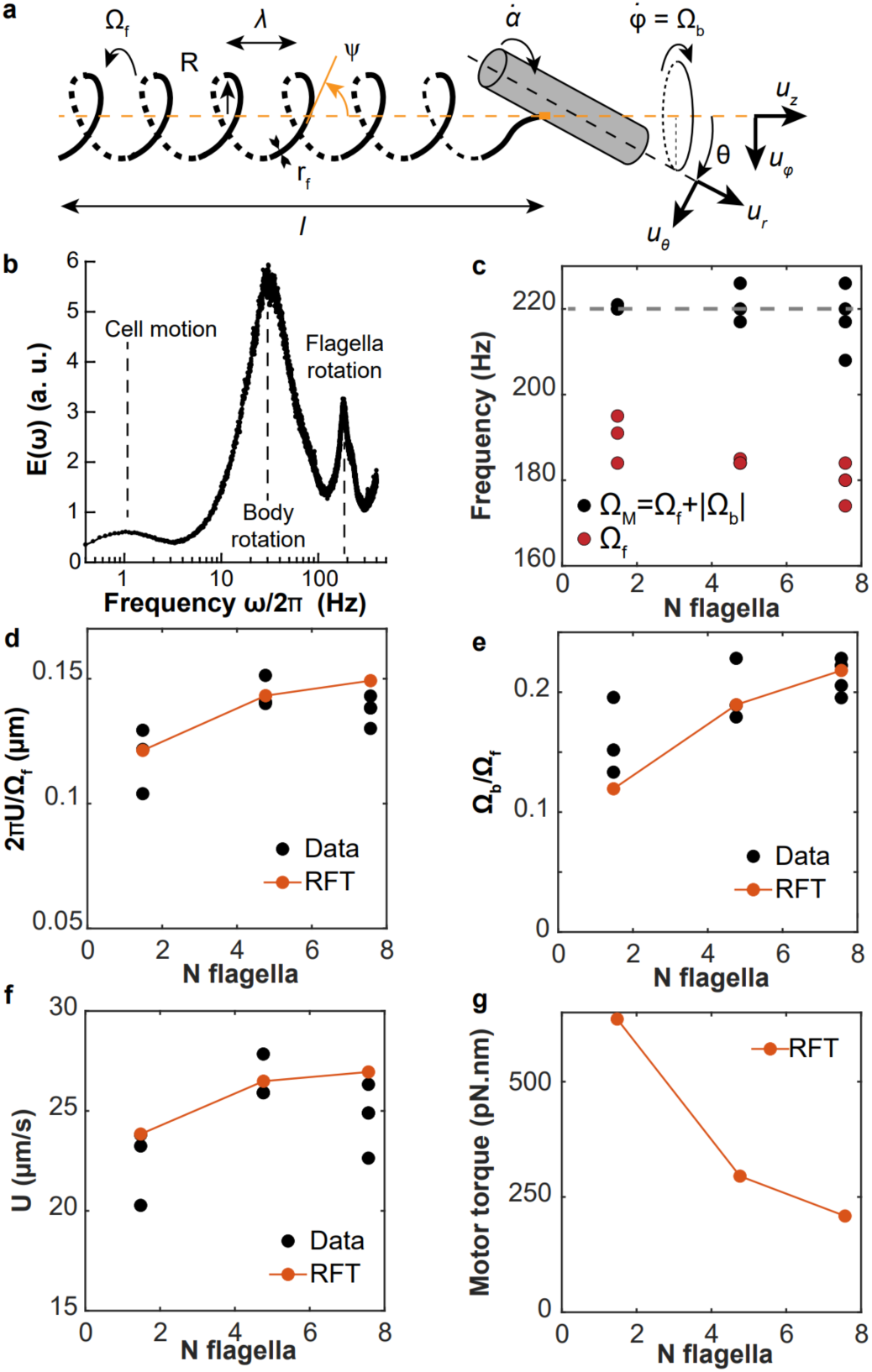
Model of flagellar propulsion. **a**, Scheme of the model of flagellar propulsion, featuring a tightly wrapped bundle rotated by N flagellar motors and a counter-rotating cell body. Geometric parameters as well as swimming speed (U), and body (Ω_!_) and flagellar (Ω”) rotation speeds are indicated. **b**, Example of normalized power spectrum *E*(ω) obtained by dark field flicker microscopy (DFFM) for MG1655 *WT* cells. The second and third peaks measure the rotation frequencies of the cell body and flagellum, respectively. **c**, Measured rotation frequencies. The black points are motor frequencies (sum of flagellum and cell body rotation rates), and the red points are flagellar frequencies. The gray dotted line is the motor frequency used in the model (220 Hz). **d-f**, Resistive force theory predictions (RFT, Supplementary Note 2, Eqs. 2.8 (**d**), 2.9 (**e**), and 2.5-2.6 (**f**)) compared to experimental measurements (Data) for *Ptac0*, MG1655 *WT* and *Ptac50* strains (from left to right) using DDM and DFFM (see Supplementary Note 1). In **f**, the motor rotation speed is set to 220 Hz, consistent with the experimental value found in **c**. **g**, Predicted flagellar motor torque (Supplementary Note 2, Eq. 2.11) for the three same strains using RFT. (**c-g**) The strains are indexed by the measured average number of flagella they harbor. (**c-g**) Each experimental data point is a biological replicate.

**Extended Data Fig. 6.**
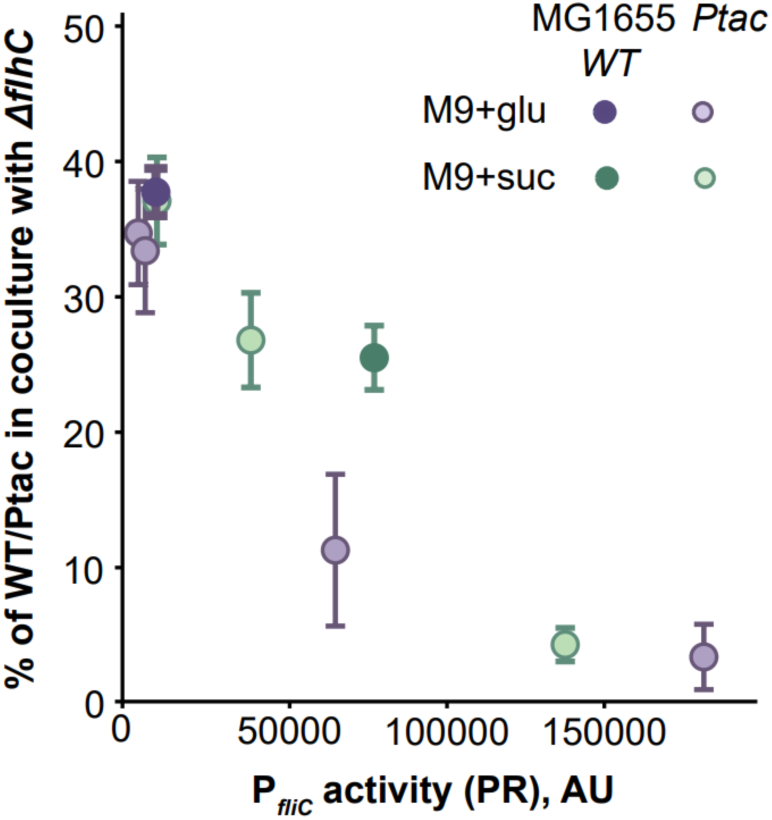
Growth fitness cost of flagellar gene expression in minimal medium. Strains were initially co-inoculated in a 1:1 ratio, and fitness cost was determined as the percentage of either MG1655 *WT* or *Ptac* strain (induced by different concentrations of IPTG) in the co-cultures with the non-flagellated *ΔflhC* strain after 72 h of incubation with shaking (200 rpm). P*_fliC_* activity measured in the plate reader (PR) was used to plot the data. The mean ± s.d. values (*n* = 3 biological replicates) are shown.

**Extended Data Fig. 7.**
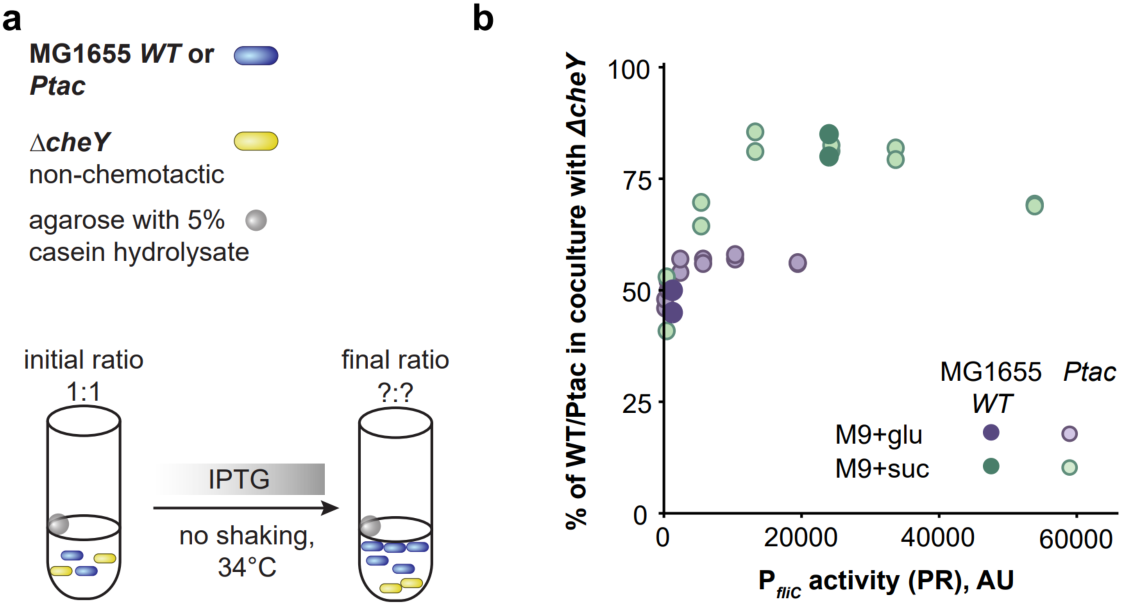
Growth fitness benefit of flagellar gene expression in minimal medium. Schematic overview (**a**) and results (**b**) of pairwise growth competition between chemotactic MG1655 *WT* or *Ptac* strain (induced by different concentrations of IPTG) and non-chemotactic *ΔcheY* grown in the presence of localized nutrient source (agarose beads containing 5% casein hydrolysate) for 72 h without shaking. Strains were initially co-inoculated in a 1:1 ratio, and fitness benefit was quantified as the percentage of MG1655 *WT* or *Ptac* strain in the mixed population at the end of the experiment (*n* = 2 biological replicates). P*_fliC_* activity measured in the plate reader (PR) was used to plot the data (*n* = 1). Note that a different plate reader was used in these experiments, for consistency with a previous publication^21^ (see Methods).

**Extended Data Fig. 8.**
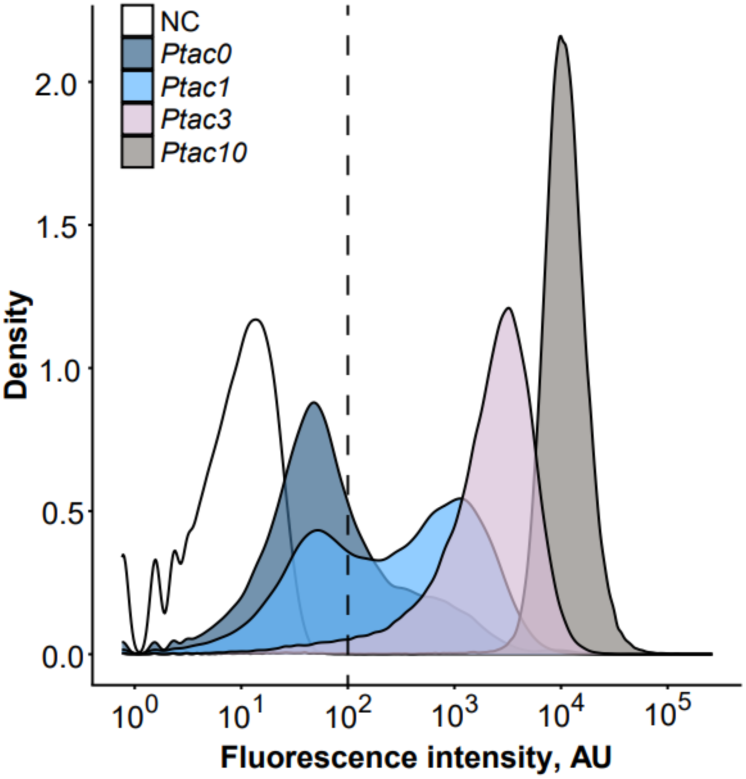
Flow cytometry measurements of P*_fliC_*-GFP reporter activity in the *Ptac* strain population grown in M9 succinate. Flagellar gene expression was induced by different concentrations of IPTG (indicated by numbers); the *Ptac* strain lacking the reporter plasmid served as negative control (NC). The vertical dashed line indicates the gate used to distinguish GFP-positive from GFP-negative cells.

**Extended Data Fig. 9.**
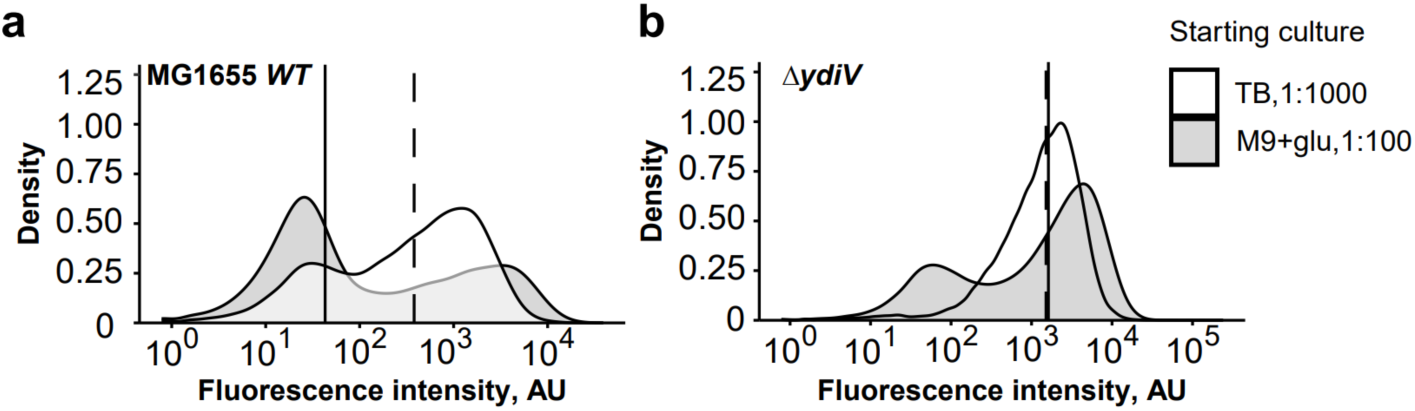
Flow cytometry analysis of P*_fliC_*-GFP reporter activity in the populations of MG1655 *WT* and *ΔydiV* strains. Cells were subjected to prolonged incubation under catabolite repression in M9 glucose either by using a higher dilution of the overnight culture (TB, 1:1000) or by pre-growing the overnight culture in M9 glucose (M9+glu, 1:100). The vertical line indicates median P*_fliC_* activity (solid for M9+glu, 1:100; dashed for TB, 1:1000). Representative data from one replicate are shown.

**Extended Data Fig. 10.**
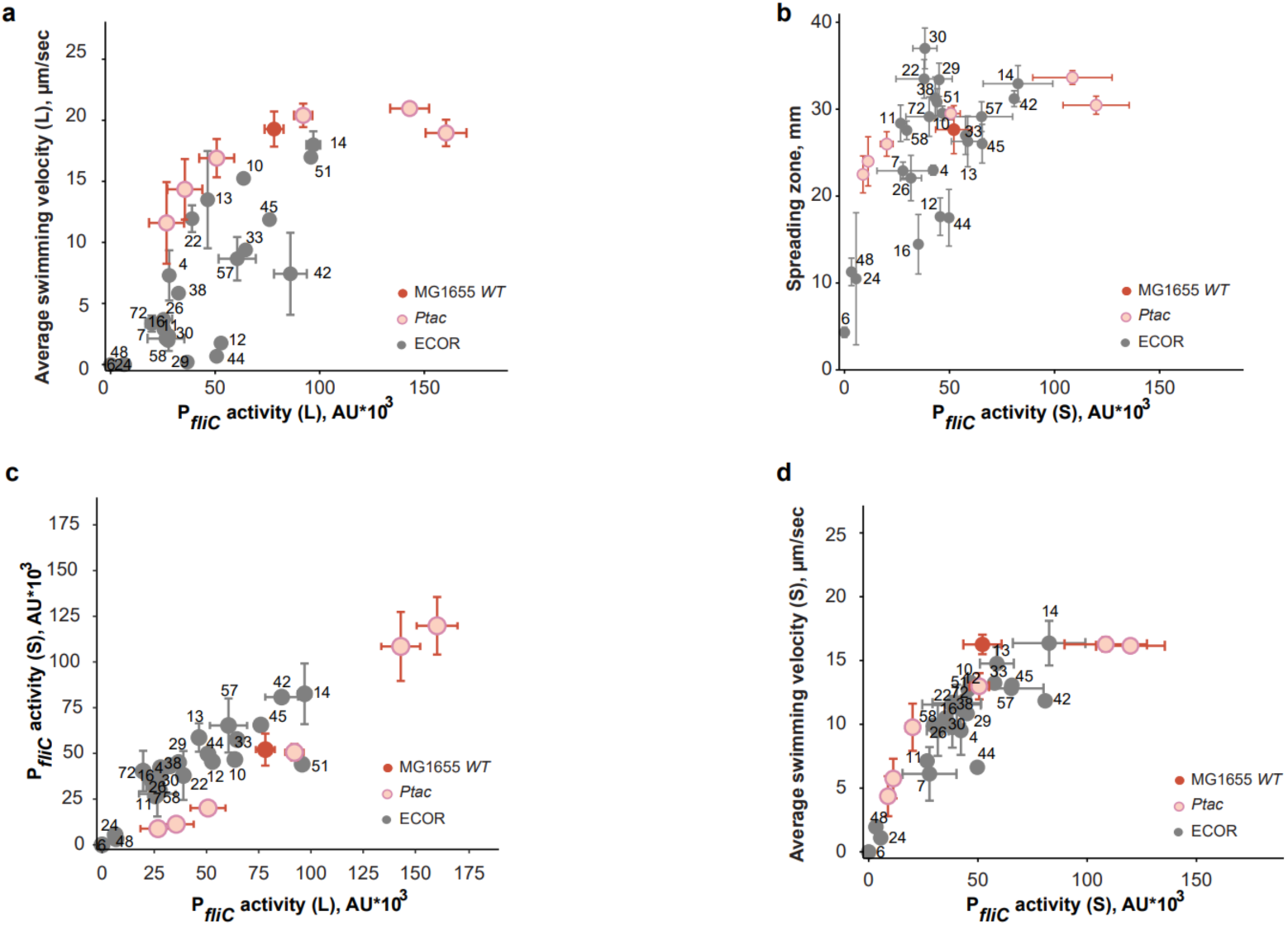
Relations between P*_fliC_* reporter activity, swimming and spreading for MG1655 *WT*, *Ptac* and the cohort of 24 natural *E. coli* isolates. Panels (**a**, **c**-**d**) contain the same data as panels (**a**, **c**-**d**) in Fig. 4 (*n =* 3 biological replicates for 10 ECOR strains, mean ± s.d.) supplemented by the non-replicate measurements for the remaining 14 strains. **b**, Dependence of spreading zone diameter (in mm) in porous 0.27% TB agar (data from Fig. 4b) on P*_fliC_* reporter activity in *E. coli* strains grown on semi-solid (S) 0.5% TB agar (data from (**c**)).

