## Supplementary Information for "Physics and physiology determine strategies of bacterial investment in flagellar motility"

### Supplementary Note 1

#### Differential Dynamic Microscopy

The principles of differential dynamic microscopy (DDM) and its application to measuring bacterial swimming behavior have been already described in detail[1-4]. Thus, we only briefly summarize the main points here.

The method exploits the rate of changes in pixel intensity in an image being proportional to the speed of motion of the object to measure it. The analysis is carried out in spatial Fourier space because the structure of the information is more easily exploitable in this format. We can write the pixel intensity of a movie frame featuring $N$ moving bacteria in a static background $I_{bg}$ as:

$$I(\boldsymbol{x},t) = I_{bg}(\boldsymbol{x}) + \sum_{i=0}^{N} I_p\big(z_i(t), \boldsymbol{x} - \boldsymbol{x}_i(t)\big) + \epsilon(\boldsymbol{x},t) \tag{1.1}$$

Where $\boldsymbol{x}_i(t)$ is the 2D position and $z_i(t)$ the position relative to the focal plane of bacterium $i$, $\epsilon(\boldsymbol{x},t)$ is the camera shot noise, and $I_p$ is the pixelated point spread function of the bacterium, that depends on the optics of the microscope and the camera and is centered around the position of the bacterium $\boldsymbol{x}_i(t)$. DDM first computes the spatial Fourier transform of this intensity:

$$I(\boldsymbol{q},t) = \int d\boldsymbol{x}\, e^{-\boldsymbol{q}\cdot\boldsymbol{x}}\, I(\boldsymbol{x},t) \; = I_{bg}(\boldsymbol{q}) + \epsilon(\boldsymbol{q},t) + \sum_{i=0}^{N} e^{-\boldsymbol{q}\cdot\boldsymbol{x}_i(t)}\, I_p(z_i(t), \boldsymbol{q}) \tag{1.2}$$

And then focusses on the differential intensity correlation function (DCIF):

$$D(\boldsymbol{q},dt) = \langle |I(\boldsymbol{q},t+dt) - I(\boldsymbol{q},t)|^2 \rangle \tag{1.3}$$

Under the assumption that $I_p(z_i(t), \boldsymbol{q})$ evolves much more slowly than $e^{-\boldsymbol{q}\cdot\boldsymbol{x}_i(t)}$, which has been verified for the low magnification optics we use and in the range of wave numbers $q$ we consider[2,3]. In the case of a dilute system where the motion of different bacteria can be seen as independent from each other, we can write the DCIF as[1,2]:

$$D(\boldsymbol{q},dt) = 2N\left|\bar{I}_p(\boldsymbol{q})\right|^2 S(\boldsymbol{q})\big(1 - f(\boldsymbol{q},dt)\big) + \langle |\epsilon(q)|^2 \rangle \tag{1.4}$$

With the structure factor $S(\boldsymbol{q}) = \langle e^{-\boldsymbol{q}\cdot(x_i - x_j)} \rangle_{i,j}$ that quantifies the spatial structure of the cell suspension, $\bar{I}_p(\boldsymbol{q})$ averaging over visible cells at different $z$, and the intermediate scattering function (ISF):

$$f(\boldsymbol{q},dt) = \langle e^{-\boldsymbol{q}\cdot\Delta x(dt)} \rangle = \int d\boldsymbol{x}\, e^{-\boldsymbol{q}\cdot\boldsymbol{x}}\, p(\boldsymbol{x}|dt) \tag{1.5}$$

Which is the Fourier transform of the probability of displacements $p(\boldsymbol{x}|dt)$ of the bacteria during a time step $dt$, and thus fully characterizes the dynamics of the system. The functional form of $f(\boldsymbol{q},dt)$ is known for various types of motion, including swimming cells and diffusing non-motile cells.

Since our sample contains both motile and non-motile cells, we fit the experimentally determined DCIF with the following model[2]:

$$D(\boldsymbol{q},dt) = A(q)\,\Big(1 - f_{ps}(\boldsymbol{q},dt)\Big) + B(q) \tag{1.6}$$

With the intermediate scattering function accounting for only a fraction $\phi_M$ of cells being motile:

$$f_{ps}(\boldsymbol{q}, dt) = \big(\phi_M f_M(\boldsymbol{q}, dt) + (1 - \phi_M)\big) f_B(\boldsymbol{q}, dt) \tag{1.7}$$

The Brownian ISF accounts for the Brownian motion of the cells (primarily the non-motile ones):

$$f_B(\boldsymbol{q}, dt) = \exp(-D_0 q^2 dt) \tag{1.8}$$

With $D_0$ their diffusion coefficient. This function decays from 1 to 0 in a typical time $dt \sim 1/D_0 q^2$.

The ISF of the motile cells has the form:

$$f_M(\boldsymbol{q}, dt) = g(q v_0 dt \mid \sigma_v / v_0) \tag{1.9}$$

With $v_0$ the average swimming speed of the population and $\sigma_v$ its variance. In the limit $\sigma_v = 0$, the function $g(x)$ is a cardinal sine ($\sin x/x$). In Wilson *et al* (2011)[2], it was computed analytically in the case of Schultz distributed velocities, a model that proved proficient:

$$g(x \mid y) = \left(\frac{Z+1}{Zx}\right) \frac{\sin\left(Z \tan^{-1}\left(\frac{x}{Z+1}\right)\right)}{\left(1 + \left(\frac{x}{Z+1}\right)^2\right)^{\frac{Z}{2}}} \tag{1.10}$$

With $Z = (y^2 - 1)/y^2$. In all cases, the function decays from 1 to 0 on a typical time $dt \sim 1/q v_0$.

In practice, the DCIF $D(\boldsymbol{q}, dt)$, which is computed over the accessible range of q given our pixel size and the width of our camera field of view, displays two distinct increases as a function of $dt$ (Extended Data Fig. 2a). These are respectively due to the motile cells (corresponding to the $1/q v_0$ decay timescale of $f_{ps}$) and the non-motile ones ($1/D_0 q^2$ timescale). The relative amplitudes of the two increases indicate the fraction of cells that are swimming (Extended Data Fig. 2a). Since the time scales have different dependences in $q$, we independently fit $D(\boldsymbol{q}, dt)$ as a function of $dt$ by the model (Eqs. 1.6-1.10) for different *q*, and verify the consistency of the fit parameters as a function of *q* (Extended Data Fig. 2b). We thus are able to validate the model and in particular the motile and Brownian nature of the motion of the two subpopulations. For very small and very large $q$, the fits fail, because the correlation function does not fully decorrelate over the duration of the experiment (small $q$) or because the signal over noise ratio is too small (large $q$). We thus reject the resulting very noisy values for the fitted parameters. We then use the average value of the parameters over the valid $q$ range as the measured values for the given experiment, while the standard deviation measures the accuracy of our estimation of said parameters.

**Dark Field Flicker Microscopy**

Dark field flicker microscopy (DFFM) is a recently developed optical microscopy method, which exploits the specificity of dark field illumination to measure the rotation speed of the cell body and the flagellum[5,6]. Under dark field illumination, the objective collects only the light that is scattered by the microscopic objects that are in focus (Extended Data Fig. 5b). In the regime of Mie scattering in which bacteria lie, the directions in which an *anisotropic* object (like the cell body or the flagellum) scatters incident light depends on the angle between the object main axis and the incident light. The rotation of this axis thus modifies the main scattering directions and therefore the amount of light collected by the objective: the image of the object flickers at a frequency that is equal to the rotation frequency of the object. This flickering can thus be exploited to measure said frequency.

In practice, a bacterial suspension prepared as for DDM ($OD_{600}$ ~ 0.1-0.2) is observed at 10x magnification (NA=0.3) under dark field illumination, and a 512 x 512 $px^2$ (1 px = 0.7 µm) field of view is recorded at 800 frames/s during $10^4$ frames with an EoSens 4CPX CMOS camera, far from the sample surfaces. The movie is divided in independent 8x8 $px^2$ submovies, in which there is on average no more than one cell at a time. The temporal power spectrum of the average intensity of the submovies is computed as

$$S(\omega) = \langle |\overline{I_k}(\omega)|^2 \rangle_k \tag{1.11}$$

With $\overline{I_k}(\omega)$ the Fourier transform of the average intensity of submovie k:

$$\overline{I_k}(\omega) = \int dt\, e^{-i\omega t} \langle I(x,t) \rangle_{x \in V(k)} \tag{1.12}$$

The power spectrum is then corrected for the effects of the cells Brownian motion by computing:

$$E(\omega) = \omega^2 S(\omega) \tag{1.13}$$

For swimming *E. coli*, the corrected power spectrum $E(\omega)$ displays three characteristic peaks above a flat background (Extended Data Fig. 5 c). The first one, at a frequency of about 1 Hz corresponds to cells moving in and out of the submovie boxes. The second one, at about 20-40 Hz corresponds to the rotation rate of the cell body $\Omega_b$, while the third, at about 200 Hz, corresponds to the rotation rate of the flagellum. The location of the maxima of the peaks is estimated via local Gaussian fits and used as a measure of the average rotation speeds within the sample.

### Supplementary Note 2

#### Detailed description of RFT model

The model extends on classical force balance analysis of mono-flagellated propulsion [7,8], and accounts for our measurements of swimming speeds as well as cell body and flagellar rotation frequencies. A schematic of the model for flagellar rotation is provided in Extended Data Fig. 5a. We assume that the *N* flagella of the cell bundle together tightly, and that the N motors are aligned along the axis of the bundle. Hence, the bundle is described as a helix of increased thickness $r_f = r\ N^{1/2}$. This assumption is based on previous experiments and simulations showing that the fluid flow generated by two helices is equivalent to the fluid flow generated by one helix with a thicker radius [9], and that the thrust generated by a bundle of helices doesn't depend much on the bundle configuration [10]. We also account for the increased length of the flagella at higher induction of *flhDC*.

Since no net torque or force are applied on the system, the friction forces and torques applying on the flagellar bundle $(F_f, \Gamma_f)$ and the cell body $(F_b, \Gamma_b)$ equilibrate in the low Reynolds number fluid:

$$\begin{gathered} F_f + F_b = 0 \\ \Gamma_f + \Gamma_b = 0 \end{gathered} \tag{2.1}$$

The friction forces and torques on the flagellar bundle, modelled as a helix, relate to its rotation speed $\Omega_f$ and the free-swimming speed $U$ of the cell via

$$\begin{pmatrix} F_f \\ \Gamma_f \end{pmatrix} = \begin{pmatrix} \mu_{TT}^f & -\mu_{TR}^f \\ -\mu_{TR}^f & \mu_{RR}^f \end{pmatrix} \begin{pmatrix} U \\ \Omega_{\mathrm{f}} \end{pmatrix} \tag{2.2}$$

Where $\boldsymbol{\mu}_X^f$ are the friction coefficient for translation (X=TT), rotation (X=RR) and rotation-translation coupling resulting from the flagellum being chiral (X=TR). We model the cell body as an (achiral) rod that wobbles. The forces and torques exerted on it relate to body rotation speed $\Omega_b$ and swimming speed $U$ via

$$\begin{pmatrix} F_b \\ \Gamma_b \end{pmatrix} = \begin{pmatrix} \mu_{T,b}\ U \\ \mu_{R,b}\ \Omega_b \end{pmatrix} \tag{2.3}$$

Where $\mu_{T,b}$ and $\mu_{R,b}$ are translational and rotational effective friction coefficients. The expressions of the coefficients as a function of the bundle and body geometric parameters are determined below (Eqs. 2.12-2.17 for the flagellum, and 2.27-2.28 and 2.30 for the cell body).

We assume that the motors all rotate at the same speed $\Omega_m$, which we assume constant based on our observations (i.e., presumably at the maximum rotation speed):

$$\mathit{\Omega}_f - \mathit{\Omega}_b = \mathit{\Omega}_m = \Omega_{Max} \tag{2.4}$$

The set of equations (2.1-2.4) solves readily and provides relations between the swimming speed and the motor rotation speed:

$$U = Z_U\ \Omega_m \tag{2.5}$$

The proportionality coefficient $Z_U$ depends on the number of flagella N and their length and reads:

$$Z_U = \frac{\mu_{TR}^f}{\mu_{t,\parallel} + \mu_{TT}^f} \frac{1}{1 + \mu_{R,\text{red}}^f / \mu_{R,\parallel}} \tag{2.6}$$

With the reduced flagellar rotational friction coefficient

$$\mu_{R,\text{red}}^f = \mu_{RR}^f - \frac{\left(\mu_{TR}^f\right)^2}{\mu_{T,\parallel} + \mu_{TT}^f} \tag{2.7}$$

Additionally, we obtain the following expressions for the swimming speed and the cell body rotation speed relative to the flagellar rotation speed:

$$\frac{U}{\Omega_f} = \frac{\mu_{TR}^f}{\mu_{T,b} + \mu_{TT}^f} \tag{2.8}$$

$$\frac{|\Omega_\text{b}|}{\Omega_f} = \frac{\mu_{R,\text{red}}^f}{\mu_{R,b}} \tag{2.9}$$

These ratios are plotted in Extended Data Fig. 5d, e for our experimental measurements compared with the resistive force theory (RFT) predictions, which account for both the length and number of flagella increasing (see below). The agreement is overall excellent. Consequently, the RFT predicts very well the swimming speed for the different strains (Extended Data Fig. 5f).

The torque generated by the motors relate to flagellar and cell body torques as:

$$\left|\Gamma_f\right| = |\Gamma_b| = N\Gamma_m \tag{2.10}$$

Which, combined with Eqs. (2.1-2.3), yields:

$$\Gamma_m = \frac{1}{N} \frac{\mu_{R,\text{red}}^f}{1 + \mu_{R,\text{red}}^f / \mu_{R,b}} \, \Omega_m \tag{2.11}$$

The predicted torque generated by each motor thus decreases as a function of N (Extended Data Fig. 5g), and never exceeds 700 pN.nm, far from the ~2500 pN.nm maximal torque that the motor can generate, which might explain why we observe a constant, presumably maximal, motor rotation speed for all strains.

Our RFT model overestimates by about 10% the experimental value of $U/\Omega_f$ and thus of $U$ for the strain with the largest number of flagella, and thus do not capture the reduction of speed there. This discrepancy is not surprising since our model becomes less realistic with increasing flagellar number, and a number of factors, including hydrodynamic effects and tangling or steric effects within the bundle could increase drag ($\mu_{TT}^f$) and/or reduce thrust ($\mu_{TR}^f$) compared to RFT prediction.

**Expression of friction coefficients**

***Flagellum***

The friction coefficients of the flagellum modeled as a helix are [8,11-13]:

$$\mu_{TT}^{f} = K_n\, l \sin\psi \left(\tan\psi + \frac{\gamma_k}{\tan\psi}\right) \tag{2.12}$$

$$\mu_{TR}^{f} = K_n\, l\, R \sin\psi \;(1-\gamma_k) \tag{2.13}$$

$$\mu_{RR}^{f} = K_n\, l\, R^2 \sin\psi \left(\frac{1}{\tan\psi} + \gamma_k \tan\psi\right) \tag{2.14}$$

With the coefficients

$$K_n = 4\pi\eta/\left(\ln\frac{c\lambda}{r_f} + 0.5\right) \tag{2.15}$$

$$\gamma_k = 0.7 \tag{2.16}$$

$$\tan\psi = 2\pi R/\lambda \tag{2.17}$$

where the helix parameters are as indicated on Extended Data Fig. 5a, and the medium viscosity is $\eta$. The length of the helix $l$ varied between 5 and 8.5 µm, according to measured values in the different strains (Fig. 2), which were least-square fitted as $l = 5.3983 + 3.6564 \log_{10} N$ to quantify the dependence of flagellar length on flagellar number. We fixed the helix wavelength $\lambda = 2.3$ µm, radius $R = 0.2$ µm, and flagellar filament thickness $r = 0.02$ µm according to previous measurements [14,15]. The bundle thickness is $r_f = r\, N^{1/2}$ to account for the increased cross-section of the bundle. The coefficient $c$ in Eq. (2.15) takes various values through the literature [8,13], from $c = 2$ in Gray and Hancock's work[11] to $c = 0.18$ in Lighthill's work[12]. We chose $c = 0.25$, which is close to Lighthill's prediction accounting for some of the hydrodynamic coupling throughout the helix [12].

***Cell body***

We model the cell body as a rod of length $L = 2.5$ µm, diameter $d = 0.75$ µm, aspect ratio $p = L/d$, consistently with the typical values for our strain. The rod is inclined with an angle $\theta$ relative to the axis of propulsion ($u_z$) and it rotates about this axis at frequency $\dot{\varphi} = \Omega_b$, which we can measure with DFFM. The cell body also rotate at a frequency $\dot{\alpha}$ about its own axis ($u_r$). Since this rotation does not affect the scattering of light by the cell body, $\dot{\alpha}$ is not detectable by DFFM. The pulsation vector accounting for these two rotations is:

$$\boldsymbol{\omega} = \sin\theta\, \dot{\varphi}\, u_z + \dot{\alpha}\, u_r \tag{2.18}$$

Projecting this vector on the axes parallel ($u_r$) and perpendicular ($u_\theta$) to the rod, we can compute the components of the torque acting on the rod:

$$\Gamma_r = \zeta_{\parallel}^{R} \boldsymbol{\omega} \cdot u_r = \zeta_{\parallel}^{R}(\sin\theta\cos\theta\, \dot{\varphi} + \dot{\alpha}) \tag{2.19}$$

$$\Gamma_\theta = \zeta_{\perp}^{R} \boldsymbol{\omega} \cdot u_\theta = \zeta_{\perp}^{R}(-(\sin\theta)^2 \dot{\varphi}) \tag{2.20}$$

The rotational friction coefficients about both axes read [16,17,]:

$$\zeta_{\perp}^{R} = \frac{\pi}{3}\eta L^3/\left(\ln p - 0.662 + 0.917/p - 0.050/p^2 + o(1/p^2)\right) \tag{2.21}$$

$$\zeta_{\parallel}^{R} = \frac{3.841\pi}{4}\eta L d^2\left(1 + 1.119\times 10^{-4} + 0.6884/p + 0.2019/p^2 + o(1/p^2)\right) \tag{2.22}$$

The torques perpendicular and parallel to the axis of propulsion of the cell are then:

$$\Gamma_z = \Gamma_r \, u_r \cdot u_z + \Gamma_\theta \, u_\theta \cdot u_z = \zeta_\perp^R (\sin\theta)^3 \dot{\varphi} + \zeta_\parallel^R (\sin\theta \, (\cos\theta)^2 \dot{\varphi} + \dot{\alpha} \cos\theta) \tag{2.23}$$

$$\Gamma_\varphi = -(\zeta_\perp^R - \zeta_\parallel^R)(\sin\theta)^2 \cos\theta \, \dot{\varphi} + \zeta_\parallel^R \sin\theta \, \dot{\alpha} \tag{2.24}$$

While the torque along z-axis is balanced by the flagellar torque, the perpendicular torque must be zero ($\Gamma_\varphi = 0$), which yields, assuming $\sin\theta \neq 0$:

$$\zeta_\parallel^R \dot{\alpha} = (\zeta_\perp^R - \zeta_\parallel^R) \sin\theta \cos\theta \, \dot{\varphi} \tag{2.25}$$

Substituting in Eq. (2.23), we obtain:

$$\Gamma_z = \zeta_\perp^R \sin\theta \, \dot{\varphi} \tag{2.26}$$

Hence the effective rotational friction coefficient is:

$$\mu_{R,b} = \zeta_\perp^R \sin\theta \tag{2.27}$$

Furthermore, we assume that the rotation of the cell body is about a fixed anchor point, which constrains $\dot{\alpha} = \dot{\varphi}$, and gives the value of the angle $\theta$ from Eq. (2.25):

$$\sin 2\theta = 2 \frac{\zeta_\parallel^R}{(\zeta_\perp^R - \zeta_\parallel^R)} \tag{2.28}$$

For our parameter values, we obtain $\theta = 21°$, which is consistent with our microscopic observations of cell swimming at high magnification.

Similarly, the friction force resulting from translation at speed U along the z-axis is given by:

$$F_z = \boldsymbol{F} \cdot u_z = u_z \cdot (\zeta_\parallel^T u_r u_r + \zeta_\perp^T u_\theta u_\theta) \cdot (U \, u_z) = \left[\zeta_\parallel^T + (\zeta_\perp^T - \zeta_\parallel^T)(\sin\theta)^2\right] U \tag{2.29}$$

Hence the effective translation friction coefficient of the cell body is:

$$\mu_{T,b} = \zeta_\parallel^T + (\zeta_\perp^T - \zeta_\parallel^T)(\sin\theta)^2 \tag{2.30}$$

with the translational friction coefficients in the direction parallel and perpendicular to the rod [17]:

$$\zeta_\parallel^T = 2\pi\eta L / (\ln p - 0.207 + 0.980/p - 0.133/p^2 + o(1/p^2)) \tag{2.31}$$

$$\zeta_\perp^T = 4\pi\eta L / (\ln p + 0.839 + 0.185/p + 0.233/p^2 + o(1/p^2)) \tag{2.32}$$

The friction force also has a component perpendicular to the axis of motion, in direction $u_\varphi$:

$$F_\varphi = \boldsymbol{F} \cdot u_\varphi = u_\varphi \cdot (\zeta_\parallel^T u_r u_r + \zeta_\perp^T u_\theta u_\theta) \cdot (U \, u_z) = \left[(\zeta_\perp^T - \zeta_\parallel^T) \sin\theta \cos\theta\right] U \tag{2.33}$$

Which however averages out over one period of cell body rotation, since $u_\varphi$ rotates. Moreover, in our case, it is negligible compared to the z-component because $\zeta_\perp^T$ is only 20% larger than $\zeta_\parallel^T$, and $\sin\theta \cos\theta \simeq 0.33$.

### Supplementary Table 1

**Strains**

| Description | Name | Source or reference | Comments |
|---|---|---|---|
| *Escherichia coli* MG1655*Δflu* | VS1671 | Ni et al, 2020. PNAS[1] | Used as the wild-type strain (*WT*) in our study. Deletion of the *flu* gene that encodes the major *E. coli* adhesin, antigen 43, prevents autoaggregation of motile planktonic cells, thus facilitating subsequent characterization of motility. |
| MG1655*Δflu*::FLP-*lacI*-$P_{tac}$-*flhDC* | VS1683 | Ni et al, 2020. PNAS[1] | Strain with the synthetic IPTG-inducible regulation of flagellar regulon (*Ptac*) |
| W3110 | VS306 | Hayashi et al, 2006. Mol Syst Biol[2] | One of the canonical K-12 derivatives |
| RP437 | VS278 | Parkinson, 1978. J. Bacteriol[3] | Canonical *E. coli* strain for chemotaxis |
| MG1655*Δflu*::FLP*ΔydiV*::FLP | IL160 | This study | Constructed via P1 transduction from KEIO collection followed by FLP recombination |
| MG1655*Δflu*::FLP*ΔflhC*::FLP | VS1674 | Ni et al, 2020. PNAS[1] | Non-motile strain used in the pairwise competition assays |
| MG1655*Δflu*::FLP*ΔcheY*::FLP | VS1672 | Ni et al, 2020. PNAS[1] | Motile and non-chemotactic strain used in the pairwise competition assays |

**Plasmids**

| | | | |
|---|---|---|---|
| $P_{fliC}$-GFP | pVS1157 | Ni et al, 2020. PNAS[1] | GFP reporter for *fliC* promoter activity based on pUA66 plasmid. sc101 Ori, $Kan^R$ |
| pTrc99a backbone | pVS2232 | Amann and Brosius, 1985. Gene[4] | Empty vector. pBR322 ori, $Amp^R$ |
| pTrc99a-CFP | pVS129 | Press et al, 2013. PLOS Genetics[5] | pTrc99a-based construct for the IPTG-inducible expression of CFP protein |
| pTrc99a-YFP | pVS132 | Oleksiuk et al, 2011. Cell | pTrc99a-based construct for the IPTG-inducible expression of YFP protein |

### Supplementary Table 2

| Strain | Host | Phylogroup | Sensitivity to kanamycin[2] | Spreading rank[3] | Isolates used for furtner experiments | Swimming (L) ≈ Swimming (S) (good/poor)[4] | Swimming (S) > swimming (L) | Expression (S) > Expression (L) |
|---|---|---|---|---|---|---|---|---|
| 1 | Human (F)[1] | A | | no | | | | |
| 2 | Human (M) | A | Kan$^R$ | | | | | |
| 3 | Dog | A | | no | | | | |
| 4 | Human(F) | A | | good | x | | x | x |
| 5 | Human (F) | A | | no | | | | |
| 6 | Human(M) | A | | poor | x | x | | |
| 7 | Orangutan | B1 | | good | x | | x | |
| 8 | Human (F) | A | Kan$^R$ | | | | | |
| 9 | Human (F) | A | | no | | | | |
| 10 | Human(F) | A | | good | x | x | | |
| 11 | Human(F), UTI | A | | good | x | | | |
| 12 | Human(F) | A | | moderate | x | | x | |
| 13 | Human(F) | A | | good | x | x | | |
| 14 | Human(F), UTI | A | | good | x | x | | |
| 15 | Human(F) | A | | no | | | | |
| 16 | Leopard | A | | moderate | x | | | |
| 17 | Pig | A | | no | | | | |
| 18 | Celebese ape | A | Kan$^R$ | | | | | |
| 19 | Celebese ape | A | Kan$^R$ | | | | | |
| 20 | Steer | A | | poor | | | | |
| 21 | Steer | A | | no | | | | |
| 22 | Steer | A | | good | x | x | | |
| 23 | Elephant | A | | no | | | | |
| 24 | Human(F) | A | | modderate | x | x | | |
| 25 | Dog | A | | no | | | | |
| 26 | Human(infant) | B1 | | good | x | | x | |
| 27 | Giraffe | B1 | | poor | | | | |
| 28 | Human(F) | B1 | | no | | | | |
| 29 | Kangaroo rat | B1 | | good | x | | | |
| 30 | Bison | B1 | | good | x | | x | |
| 31 | Leopard | D | Kan$^R$ | | | | | |
| 32 | Giraffe | B1 | | poor | | | | |
| 33 | Sheep | B1 | | good | x | | | |
| 34 | Dog | B1 | Kan$^R$ | | | | | |
| 35 | Human (M) | D | | no | | | | |
| 36 | Human (F) | D | | no | | | | |
| 37 | Marmoset | D | Kan$^R$ | | | | | |
| 38 | Human(F) | D | | good | x | | | |
| 39 | Human(F) | D | | no | | | | |
| 40 | Human(F), UTI | D | | poor | | | | |
| 41 | Human (M) | D | | poor | | | | |
| 42 | Human(M) | D | | good | x | | | |
| 43 | Human(F) | A | | moderate | | | | |
| 44 | Coguar | D | | moderate | x | | | |
| 45 | Pig | B1 | | good | x | x | | |
| 46 | Celebese ape | D | | no | | | | |
| 47 | Sheep | D | | poor | | | | |
| 48 | Human(F), UTI | D | | moderate | x | x | | |
| 49 | Human(F) | D | | moderate | | | | |
| 50 | Human(F), UTI | D | | no | | | | |
| 51 | Human(infant) | B2 | | good | x | x | | |
| 52 | Orangutan | B2 | | no | | | | |
| 53 | Human(F) | B2 | | poor | | | | |
| 54 | Human | B2 | | poor | | | | |
| 55 | Human(F) | B2 | Kan$^R$ | | | | | |
| 56 | Human(F), UTI | B2 | Kan$^R$ | | | | | |
| 57 | Gorilla | B2 | | good | x | | | |
| 58 | Lion | B1 | | good | x | | | |
| 59 | Human (M) | B2 | | no | | | | |
| 60 | Human(F), UTI | B2 | | no | | | | |
| 61 | Human(F) | B2 | Kan$^R$ | | | | | |
| 62 | Human(F), UTI | B2 | | no | | | | |
| 63 | Human(F) | B2 | | no | | | | |
| 64 | Human(F), UTI | B2 | | poor | | | | |
| 65 | Celebese ape | B2 | Kan$^R$ | | | | | |
| 66 | Celebese ape | B1 | | poor | | | | |
| 67 | Goat | B1 | | poor | | | | |
| 68 | Giraffe | B1 | | poor | | | | |
| 69 | Celebese ape | B1 | | no | | | | |
| 70 | Gorilla | A | | no | | | | |
| 71 | Human(F), UTI | A | | no | | | | |
| 72 | Human(F), UTI | A | | good | x | | x | x |

[1]:(M) - male, (F) - female, UTI - urinary tract infection. Description is taken from Ochman, H. & Selander, R.K. Standard reference strains of Escherichia coli from natural populations. Journal of Bacteriology 157, 690-693 (1984).

[2]: Kan$^R$- the strains which grew on the LB agar + Kan (50 µg/mL)

[3]: Assessed as the diameter of the spreading zone in soft TB agar (0.27% TB agar)

[4]: Swimming velocity measured after growth in liquid (L) or on the semi-solid medium (S); good - > 10 µm/sec, poor - < 10 µm/sec
